# Attentional enhancement and suppression of stimulus-synchronized BOLD oscillations

**DOI:** 10.1101/2025.01.10.632431

**Authors:** Reebal W. Rafeh, Geoffrey N. Ngo, Lyle E. Muller, Ali R. Khan, Ravi S. Menon, Marieke Mur, Taylor W. Schmitz

## Abstract

Visual cortical neurons synchronize their firing rates to periodic visual stimuli. EEG is commonly used to study directed attention by frequency-tagging brain responses to multiple stimuli oscillating at different frequencies, but is limited by its coarse spatial resolution. Here we leverage frequency-tagging fMRI (ft-fMRI) to study the influence of directed attention on the fine-grained spatiotemporal dynamics of competing stimulus-driven visual cortical oscillations. Our analysis reveals that distinct populations of visual cortical neurons exhibit in-phase (enhancing) or anti-phase (suppressive) synchronization with the oscillating stimuli. Directed attention homogeneously increases the amplitude of anti-phase BOLD oscillations across the visual hierarchy, consistent with a distributed suppressive field. In contrast, attentional modulation of in-phase BOLD oscillations increases hierarchically from V1 to hV4. The strength of anti-phase, but not in-phase, modulation predicted psychophysical correlates of attentional performance. Our results strongly corroborate the biased competition model of attention and unveil a novel BOLD correlate of attentional suppression.

## Introduction

The biased competition model of attention proposes that populations of visual cortical neurons compete for access to attentional resources through mutual suppression and enhancement of one another’s activity. When attention is directed to one of the two stimuli, the spike rates of neurons tuned to attended features are enhanced while those tuned to competing unattended features are suppressed. Since its introduction in the mid 90s, the model has received considerable support from monkey electrophysiology research. However, the mechanistic basis of biased competition has proven more challenging to study in humans. Access to both the fine-grained spatial maps of feature-tuned neuronal activity and their temporal dynamics are required to adjudicate the simultaneous competitive influences of suppression and enhancement. *In vivo* imaging methods such as electroencephalography (EEG) and magnetoencephalography (MEG) do not afford the spatiotemporal precision required to record these patterns.

In a seminal non-human primate study which combined fMRI and electrophysiology, Shmuel et al. ^1^ demonstrated that during partial visual field stimulation, blood-oxygen-level-dependent (BOLD) signals exhibit both in-phase and anti-phase relationships with visual stimulation. Anti-phase BOLD signals were localized beyond the stimulated regions of the visual cortex and colocalized with decreases in neuronal activity. This finding indicates that while one might expect BOLD signals to increase uniformly with heightened neural spike rates under sensory stimulation, there are neighboring populations of neurons where both the BOLD signal and spike rates decrease. The magnitudes of these parallel enhancing and suppressive effects were larger than those observed during stimulus-independent spontaneous activity and were negatively correlated with one another, consistent with a stimulus-driven competitive response. Although attention was not explicitly examined in these studies, the observations of stimulus-driven enhancement and suppression of spike rates, concurrent with in-phase and anti-phase BOLD responses, in spatially adjacent populations are in accord with the biased competition model of attention.

If in-phase and anti-phase BOLD signals reflect enhancement and suppression of competing visual inputs, respectively, then experimental probes of these signals under manipulations of directed attention may provide major insights into the spatiotemporal dynamics of biased competition in humans. In our prior work, we developed an fMRI frequency-tagging method inspired from EEG Steady State Visual Evoked Potential (SSVEP) work to examine the spatiotemporal dynamics of BOLD responses to two stimuli oscillating at different frequencies in different visual field locations ^2^. We provide evidence for fine-grained anatomical dissociation of BOLD signals synchronized to the driving frequency of either stimulus. Moreover, the spatial distribution of these frequency tagged populations was stable within individuals over repeated measures, consistent with a feature-tuned map of location preferences. Here we extend this fMRI platform further by examining (1) whether frequency tagged BOLD responses synchronize either in-phase or anti-phase with respect to a stimulus-driven oscillation, (2) whether these in-phase and anti-phase responses are anatomically dissociable in the visual cortex, and (3) how in-phase and anti-phase populations are modulated by covert shifts of directed attention to one of two simultaneous oscillating visual stimuli.

## Results

### fMRI frequency-tagging reveals distinct BOLD response phases in the visual cortex

We began by estimating the phase of stimulus-evoked BOLD responses using a frequency-tagging design ^2^. Participants (n=7) performed a task where they were required to distribute their attention to two checkerboard wedges presented in the upper left or right visual hemifield (counterbalanced across participants; Fig. 1a ; Supplementary Video 1). Our behavioral analyses show that participants engaged with the task and maintained central fixation throughout the experiment, indicating that different response phases are not driven by saccadic eye-movements (Supplementary Fig. 1).

**Figure 1.**
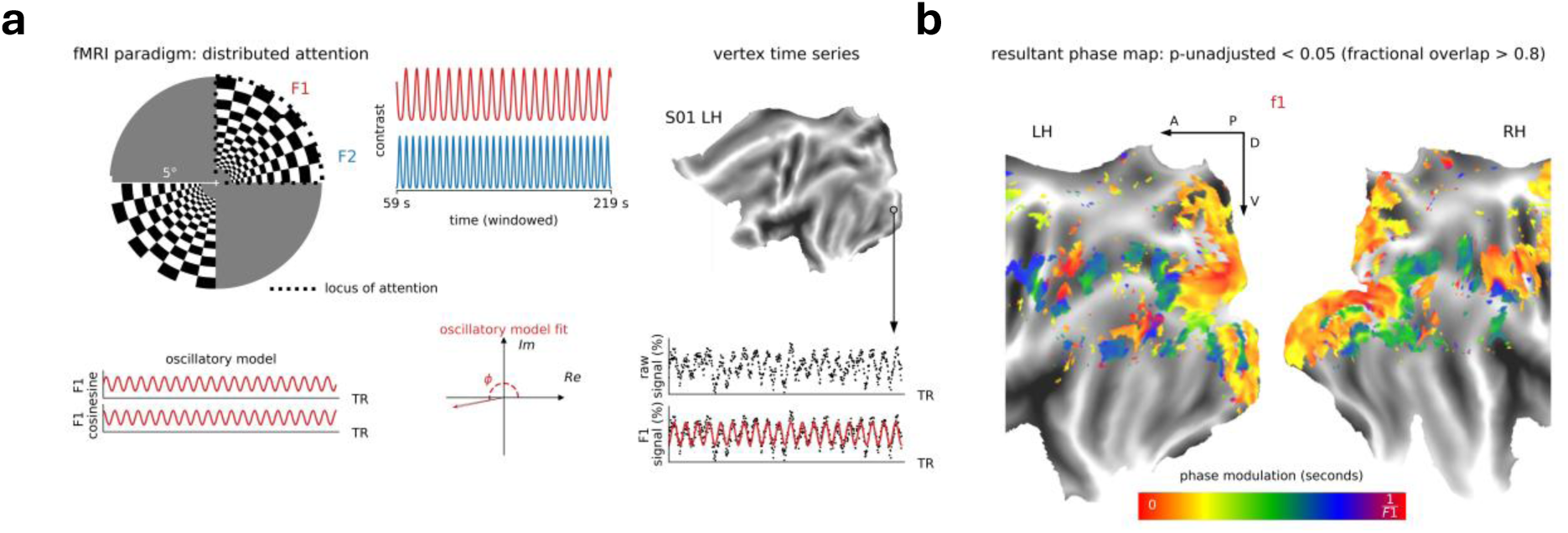
fMRI frequency-tagging reveals distinct BOLD response phases in the visual cortex. **(a)** Stimulus configuration and BOLD response phase modelling. Participants covertly directed their attention to a pair of checkerboard wedges oscillating at different frequencies (F1 = 0.125 Hz; F2 = 0.2 Hz) in an upper visual field quadrant. For each run, we analyzed the resulting vertex timeseries in a 160 s time window (59s – 219s). A sine and cosine function at each stimulation frequency was fit to the timeseries of each vertex using general linear models (GLM). The fitted sine and cosine functions were then projected onto the complex plane, providing an estimate of BOLD signal amplitude and phase. This procedure was repeated 500 times for each vertex using different resamplings of the data. **(b)** The localized visual cortical phase map at F1 projected to the flattened surface of a representative participant (S01). We thresholded vertices according to the proportion of resamples exhibiting a significant model fit at the frequency of interest (F-test; p-unadjusted < 0.05). Vertices exhibiting a significant GLM fit across at least 80% of the random samples (i.e., 400/500 resamples) were determined to be task-driven. The reported phase value for each vertex is the circular mean of phase estimates across resamples of the data. The color bar denotes the response phase scaled to the period of the examined frequency (in this case: 1/F1).

To drive neuronal oscillations at two distinct frequencies, the grating stimuli periodically alternated between periods of visibility and invisibility at either 0.125 or 0.2 Hz. Because the periodic stimuli were spatially segregated from one another, we were able to dissociate frequency-tagged neuronal oscillations in retinotopically mapped visual cortical areas which reflect preferential responses to either of the two stimuli. To access these fast neuronally-driven dynamics, we sampled fMRI at a TR = 250 ms in accordance with prior work ^3–5^ (Fig. 1a). Data were acquired from a slab covering the occipital and inferior parietal cortices.

For each participant and session, data were projected onto the cortical surface to enable vertex-wise analysis of spatiotemporal dynamics. Our frequency-tagging protocol enables convenient analysis of both the magnitude and phase of oscillatory BOLD responses. We first indexed frequency-tagged BOLD responses according to the phase of their oscillations (Fig. 1a). We observed vertices distributed throughout the primary and extrastriate visual cortex which exhibited either in-phase or anti-phase synchronization with the visual stimulation (Fig. 1b). The vertex-wise spatial profile of response phases was stable across sessions within participants (Supplementary Fig. 2a-b; surrogate map test: Kolmogorov-Smirnov p_one-tailed_ < 10^-30^, Mann-Whitney U p_one-tailed_ < 10^-20^). Moreover, we observed that the within-participant correlations of these spatial profiles were significantly higher than the between-participant correlations, suggesting that these maps reflect individually unique functional topographies (Kolmogorov-Smirnov test: p_one-tailed_ < 10^-19^, Mann-Whitney U p_one-tailed_ < 10^-15^; Supplementary Fig. 2c, 3a). These findings agree with prior work in non-human primates showing that concurrent visual cortical BOLD responses synchronize either in-phase or anti-phase with respect to a visual stimulus ^1,6^.

### Visual cortical oscillations dissociate into synchronized in-phase and anti-phase populations

Thus far, we have demonstrated that an oscillating checkerboard stimulus occupying a quadrant of the upper visual field drives a distributed pattern of frequency tagged visual and parietal cortical BOLD responses synchronized either in-phase or anti-phase to the oscillation. Because our stimulus was designed a priori to drive BOLD activity in location-tuned ventral visual areas, we focus our subsequent analyses on regions of interest (ROIs) anatomically constrained to areas V1-hV4 (see ROI definition) ^7^ (Fig. 2a).

**Figure 2.**
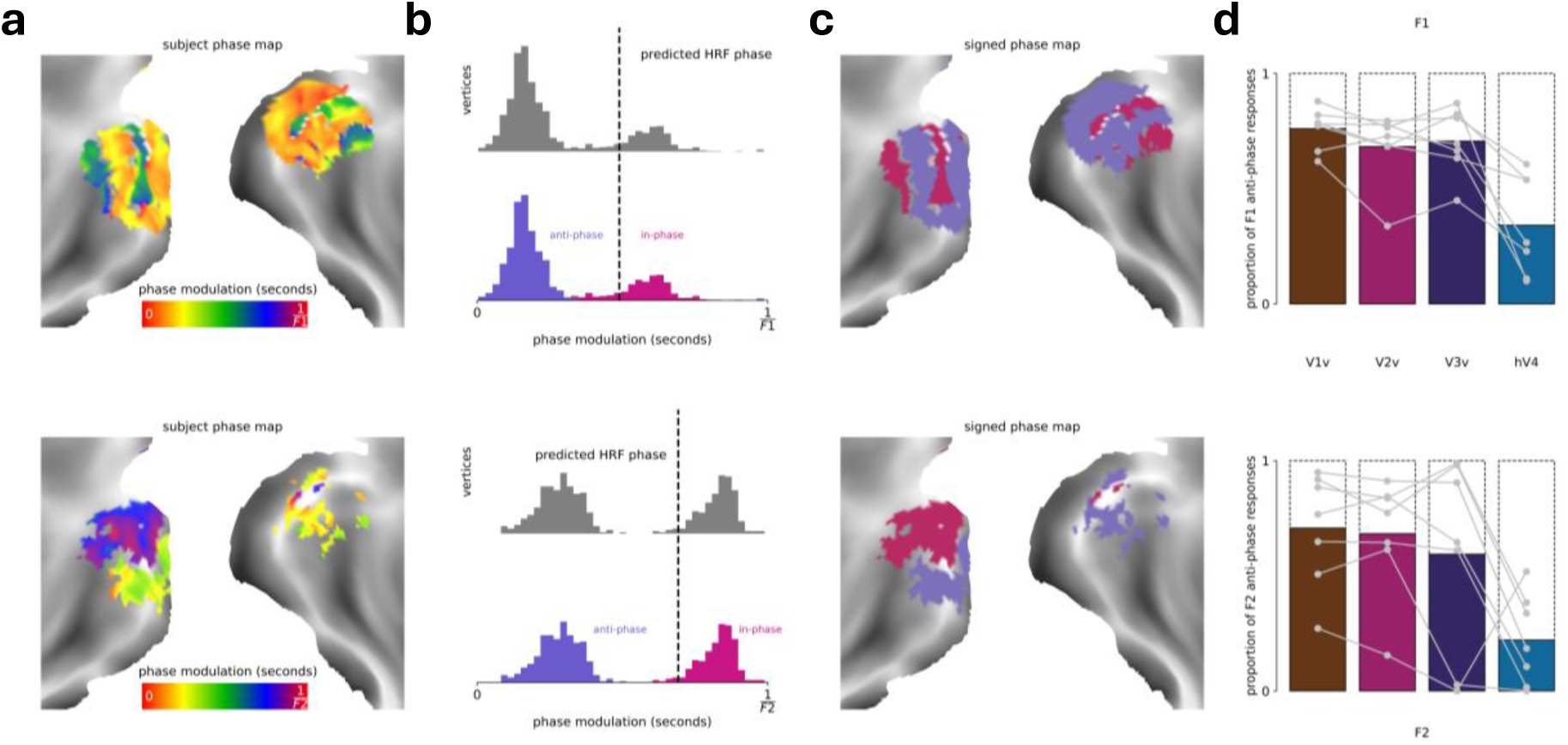
Visual cortical oscillations dissociate into synchronized in-phase and anti-phase populations. **(a)** Phase maps at each frequency of interest (F1: top; F2: bottom) were restricted to the ventral visual cortex. Color bars denote the response phase scaled to the period of the examined frequencies. **(b)** Distribution of vertex-wise response phases at each frequency. Phase data were separated into two distinct distributions by fitting a von Mises mixture model to the data. Vertices falling within the phase distribution with the smallest circular distance to the phase that would be predicted by the hemodynamic response function (HRF) were classified as in-phase, and vertices falling within the opposing distribution were classified as anti-phase. **(c)** Classified signed phase map for a representative participant. Classified phase distributions of in-phase (violet-red) and anti-phase (slate-blue) vertices shown on the cortical surface. Vertices dissociate into spatially distinct phase distributions. **(d)** Proportion of anti-phase classified vertices in different visual ROIs, expressed as ratio to the total number of vertices localized within the corresponding visual ROI. Each data point represents a single participant. Surface and distribution data in panels a-c are from a representative participant (S01).

First, we examined whether the vertices exhibiting either in-phase or anti-phase responses in V1-hV4 reflect a continuum of different phase offsets, or alternatively, whether these vertices cluster into distinct populations according to similar phase offsets. Due to the heterogeneity of BOLD response timings in the cortex, traditional fMRI modelling approaches using a canonical hemodynamic response function (HRF) may obscure variations in phase offsets of the signal ^8,9^. We address this heterogeneity by clustering responses based on their underlying phase distributions. To do so, we fit a von Mises mixture model to the vertex-wise phase offsets (Fig. 2b), which enabled us to examine if the phase offsets of vertex responses cluster into unique distributions. We examined the model fits using the Bayes Information Criterion (BIC) with increasing numbers of phase clusters. Two clusters of vertices maximized the amount of information described by the model, consistent with two temporally opposing populations (Supplementary Fig. 4a).

We next determined which of these two populations oscillates either in-phase or anti-phase with respect to the stimulus and then compared their spatial distributions across the ventral visual hierarchy. This was accomplished by convolving the stimulus time series with a gamma function to estimate the phase of a canonical in-phase response, and then by classifying each vertex according to the circular distance of its phase information from this canonical reference. We described vertices falling in the distribution closest to the predicted phase as responding in-phase, and those falling within the opposing phase distribution as anti-phase (Fig. 2c, Supplementary Fig. 3b, 4b, Supplementary Video 2). We found that anti-phase responses constituted the majority of responses in earlier visual regions (V1-V3) and the minority of responses in visual region hV4 (Fig. 2d). Moving up the cortical hierarchy, the ratio of in-phase to anti-phase responses reversed, with a significant decrease in anti-phase responses in hV4 relative to earlier visual areas in vertices responsive to F1 (Kruskal-Wallis test: H(3) = 13.5; p = 0.004), and F2 (Kruskal-Wallis test: H(3)

= 8.43; p = 0.038) (Supplementary Table 1). Consistent with a fine-grained topographic map of two opposing populations across V1-hV4, the assignment of vertices into spatially dissociated in-phase and anti-phase clusters was stable within participants over resamplings of their data (Supplementary Fig. 3b, 5). Furthermore, the vertex-wise assignment as either in-phase or anti-phase was reliable across sessions within participants (Supplementary Fig. 6a-b, in-phase Dice’s coefficient > 0.59, surrogate map test: Kolmogorov-Smirnov p_one-tailed_ < 10^-38^, Mann-Whitney U p_one-tailed_ < 10^-26^; Supplementary Fig. 6d-e; anti-phase Dice’s coefficient > 0.78, surrogate map test: Kolmogorov-Smirnov p_one-tailed_ < 10^-60^, Mann-Whitney U p_one-tailed_ < 10^-28^). Moreover, the within-participant overlap of these spatial profiles were significantly different from the between-participant overlap, suggesting that these maps reflect individually unique functional topographies of feature-tuned competitive responses (Supplementary Fig. 6c, in-phase Kolmogorov-Smirnov test: p_one-tailed_ < 10^-12^, Mann-Whitney U p_one-tailed_ < 10^-14^; Supplementary Fig. 6f, in-phase Kolmogorov-Smirnov test: p_one-tailed_ < 10^-20^, Mann-Whitney U p_one-tailed_ < 10^-22^).

### Attentional modulation of frequency-tagged BOLD oscillations in the visual cortex

We next introduced an explicit manipulation of directed attention to examine how shifts in an observer’s internal state modulate competitive dynamics in the human visual cortex. During our localizer experiment, participants distributed their attention to both of the oscillating checkerboard wedges, thereby minimizing attentional bias to either of the two frequency-tagged inputs. In the directed attention experiment, participants covertly directed their spatial attention to one of the two oscillating checkerboard wedges, thereby biasing competition to favor one of the two frequency-tagged inputs, while holding bottom-up visual stimulation identical to the localizer. This yielded two distinct attention conditions: (1) F1 attended and F2 unattended (aF1/uF2); (2) F2 attended and F1 unattended (aF2/uF1). Our primary comparisons of directed attention then focused on comparisons of BOLD oscillation amplitudes (1) between aF1 and uF1 (aF1-uF1), and (2) between aF2 and uF2 (aF2-uF2; Fig. 3; Supplementary Fig. 7a). Vertex-wise paired samples t-tests for (aF1 vs. uF1) and (aF2 vs. uF2) revealed that directed attention to either F1 or F2 significantly modulated the amplitude of BOLD oscillations in spatially dissociated vertices (p_two-tailed_<0.05, 1000 permutations; Supplementary Fig. 7b).

**Figure 3.**
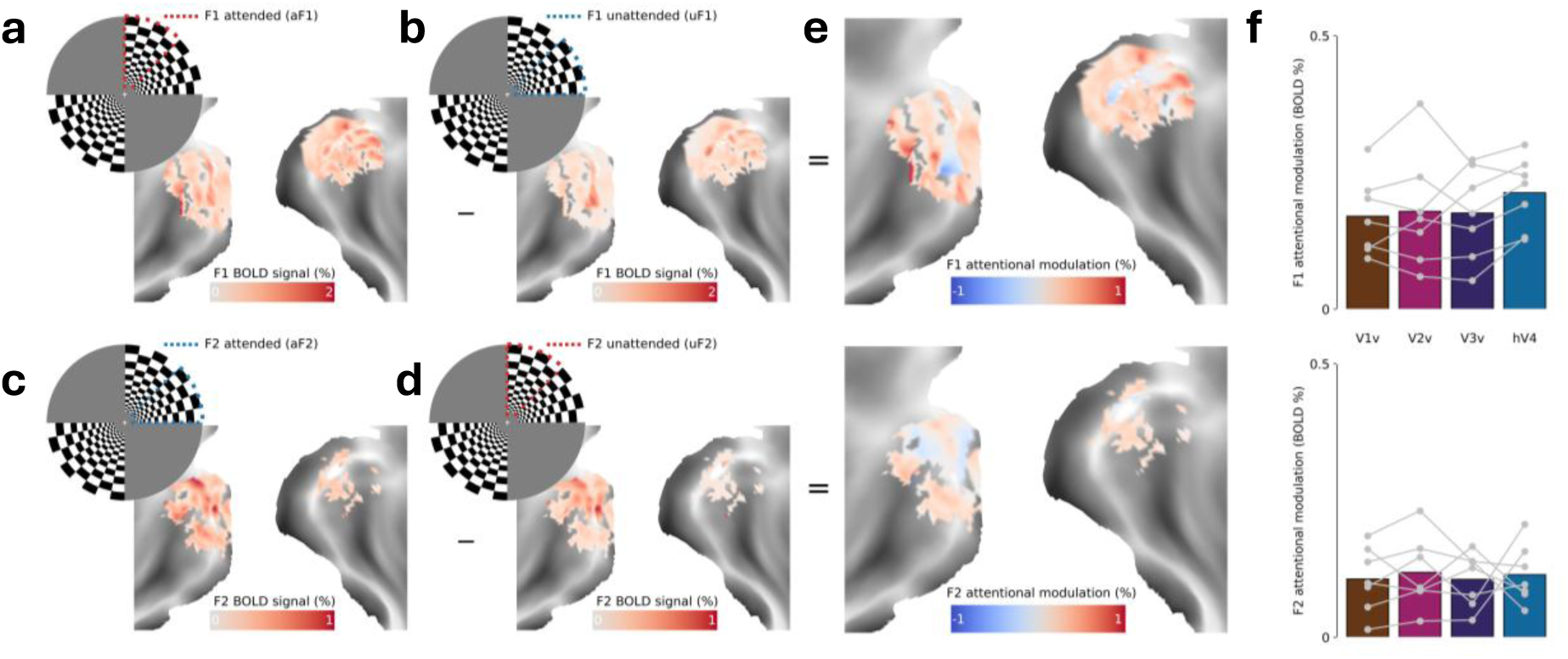
Attentional modulation of frequency-tagged BOLD oscillations in the visual cortex. F1 BOLD oscillation amplitudes during **(a)** attended F1, **(b)** unattended F1. F2 BOLD oscillation amplitudes during **(c)** attended F2, **(d)** unattended F2. **(e)** Attentional modulation map at F1 (top; aF1-uF1) and F2 (bottom; aF2-uF2). **(f)** Average attentional modulation of the amplitude of the oscillatory BOLD signal across localized vertices in different visual ROIs. Each data point represents a single participant. Surface data in panels a-e are from a representative participant (S01).

Consistent with a fine-grained map of modulation by directed attention, these vertex-wise spatial topographies of attentional modulation under aF1 and aF2 were stable across sessions (Supplementary Fig. 8a,b; surrogate map test: Kolmogorov-Smirnov p_one-tailed_ < 10^-25^, Mann-Whitney U p_one-tailed_ < 10^-19^). Furthermore, we observed that the within-participant session correlations of these spatial profiles were significantly higher than the between-participant correlations, suggesting that these maps reflect individually unique functional topographies (Kolmogorov-Smirnov test: p_one-tailed_<10^-10^, Mann-Whitney U p_one-tailed_ < 10^-12^; Supplementary Fig. 7a, 8c).

Prior human fMRI work suggests that the strength of attentional modulation of visual cortical BOLD responses to similar types of checkerboard stimuli increases moving up the visual cortical hierarchy, e.g. from V1 to TEO ^10,11^. To examine if this pattern was expressed in the spatiotemporal dynamics of frequency tagged visual cortical BOLD oscillations, we compared the strength of attentional modulation across the ROIs spanning V1 to hV4 (Fig. 3c). Although there were qualitative increases in the strength of attentional modulation in hV4 compared to earlier visual areas (V1-V3) for the aF1 condition, these effects were non-significant (Kruskal-Wallis test: H(3) = 1.65; p = 0.65), and absent altogether in aF2 (Kruskal-Wallis test: H(3) = 0.15; p = 0.98).

### Attention differentially modulates in-phase and anti-phase frequency-tagged BOLD oscillations

Thus far, we have shown that distributed attention to two wedge stimuli oscillating at different frequencies evoked concurrent in-phase and anti-phase frequency tagged BOLD oscillations (Fig. 1a,b). The visual cortical populations exhibiting in-phase and anti-phase oscillations dissociate into reliable fine-grained maps which change in their relative proportion across the visual cortical hierarchy (Fig. 2c). Moreover, we have shown that directed attention to one of these two wedge stimuli evokes strong and spatially reliable modulatory influences on the amplitudes of frequency tagged in-phase and anti-phase BOLD oscillations (Fig. 3a,b). However, the strength of this modulatory effect did not appear to change substantially as a function of vertex location along the visual cortical hierarchy (Fig. 3c). To examine if attention differentially modulates in-phase and anti-phase BOLD oscillations along the visual cortical hierarchy, we next integrated the distributed attention and directed attention datasets. Specifically, we examined how the directed attention conditions aF1 and aF2 (Fig. 3) influence oscillatory BOLD dynamics within the spatially dissociated in-phase and anti-phase vertices defined independently by the distributed attention task (under identical visual stimulation; Fig. 2).

We first examined the effect of attentional modulation on in-phase and anti-phase vertices separately for F1 (aF1 - uF1; Fig. 4a) and F2 (aF2 - uF2; Fig. 4b). For both frequencies, we found that attention significantly enhanced the amplitude of both the in-phase (Wilcoxon signed-rank test: p_F1 two-tailed_ = 0.0313, D_F1_ = 1.308; p_F2 two-tailed_ = 0.0313, D_F2_ = 1.378) and the anti-phase vertices (Wilcoxon signed-rank test: p_F1 two-tailed_ = 0.016, D_F1_ = 3.854; p_F2 two-tailed_ = 0.016, D_F2_ = 5.190). We also directly compared the magnitude of attentional modulation between the in-phase and anti-phase vertices for F1 [(aF1_anti-phase_ - uF1_anti-phase_) - (aF1_in-phase_ - uF1_in-phase_)] and F2 [(aF2_anti-phase_ - uF2_anti-phase_) - (aF2_in-phase_ - uF2_in-phase_)]. For both frequencies F1 and F2, we observed that the amplitude of BOLD oscillations was more strongly modulated in anti-phase relative to in-phase vertices (Wilcoxon signed-rank test: p_F1 two-tailed_ = 0.016, D_F1_ = 2.497; p_F2 two-tailed_ = 0.016, D_F2_ = 2.761).

**Figure 4.**
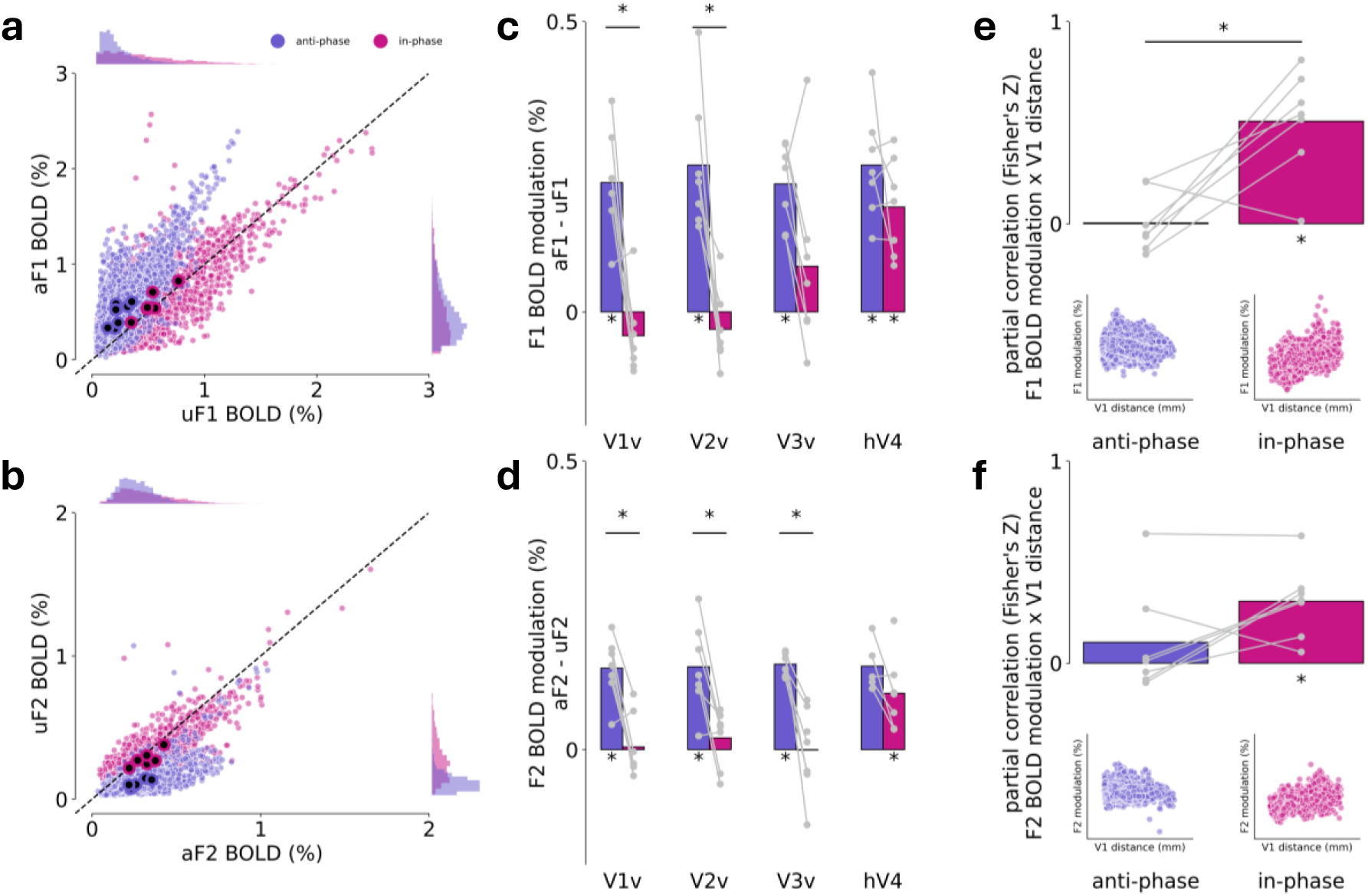
Attention differentially modulates in-phase and anti-phase frequency-tagged BOLD oscillations. **(a)** The relationship of aF1 (y-axis) to uF1 (x-axis) across all vertices of all participants for the in-phase (violet-red) and anti-phase (slate-blue) vertices (small closed circles), and the vertex-wise averages for each participant (closed large circles). Dots along the diagonal indicate no difference in BOLD modulation between aF1 and uF1. **(b)** The relationship of aF2 (x-axis) to uF2 (y-axis). Representation of the data follows the same convention as in (a). **(c)** Attention-driven modulation of the amplitudes of in-phase and anti-phase BOLD oscillations for F1 (aF1-uF1). Gray dots represent the average attentional modulation across in-phase or anti-phase vertices within an ROI for each participant. **(d)** Attention-driven modulation of the amplitudes of in-phase and anti-phase BOLD oscillations for F2 (aF2-uF2). Representation of the data follows the same convention as in (c). **(e)** and **(f)** Increasing geodesic distance from V1 relates to stronger attentional modulation of (e) the F1 amplitude and (f) the F2 amplitude of the in-phase, but not anti-phase, oscillatory BOLD signals. Insets: scatterplots showing the relationship between geodesic distance from V1 and attentional BOLD modulation for all participants’ anti-phase (left) and in-phase (right) vertices. Statistical significance was determined using a Wilcoxon signed-rank test (*P_two-tailed_ < 0.05).

Although the effects of attentional modulation on frequency tagged BOLD oscillations appear stronger overall on the vertices with anti-phase synchronization to the stimulus, it is possible this difference masks an additional interdependence, namely how attention influences in-phase and anti-phase BOLD oscillations as a function of processing stage along the visual cortical hierarchy. To examine this interdependence explicitly, we repeated the comparisons of attentional modulation on in-phase and anti-phase vertices, focusing first on F1 (aF1 - uF1; Fig. 4a) and F2 (aF2 - uF2; Fig. 4b), but now separated according to visual cortical ROI (V1-hV4; Supplementary Fig. 9). This was accomplished using four separate Kruskal-Wallis ANOVAs, one for each combination of frequency and oscillatory phase (F1 in-phase; F2 in-phase; F1 anti-phase; F2 anti-phase), modeling the effect of ROI (n=4) on attentional modulation. We found that there is a significant effect of ROI on attentional modulation for in-phase vertices at both F1 and F2 (Kruskal-Wallis test: H(3)_F1_ = 13.4, p_F1 two-tailed_ = 0.004; H(3)_F2_ = 7.61, p_F2 two-tailed_ = 0.055). By contrast, the effect of ROI was not significant for the anti-phase vertices at either F1 or F2 (Kruskal-Wallis test: H(3)_F1_ = 0.36, p_F1 two-tailed_ = 0.949; H(3)p_F2_ = 0.28, p_F2 two-tailed_ = 0.965). Post hoc comparisons evaluating the effects of attentional modulation on in-phase and anti-phase oscillatory BOLD amplitudes within each ROI are displayed in Fig. 4c,d and Supplementary Table 2. Overall, a clear pattern emerges where directed attention differentially modulates the amplitude of in-phase BOLD oscillations moving up the visual hierarchy, with the strongest effects observed in hV4, whereas attentional modulation of anti-phase BOLD oscillations does not change with hierarchical stage.

We next examined if ROI differentiated the effects of attentional modulation in the direct comparison between the in-phase and anti-phase vertices for F1 [(aF1_anti-phase_ - uF1_anti-phase_) - (aF1_in-phase_ - uF1_in-phase_)] and F2 [(aF2_anti-phase_ - uF2_anti-phase_) - (aF2_in-phase_ - uF2_in-phase_)]. We found that there is a significant increase in attentional modulation of anti-phase relative to in-phase vertices in earlier visual regions (Wilcoxon signed-rank test V1: p_F1 two-tailed_ = 0.0313, D_F1_ = 3.254; p_F2 two-tailed_ = 0.0313, D_F2_ = 2.523; V2: p_F1 two-tailed_ = 0.016, D_F1_ = 2.945; p_F2 two-tailed_ = 0.0313, D_F2_ = 1.901; V3:p_F1 two-tailed_ = 0.078, D_F1_ = 1.156; p_F2 two-tailed_ = 0.0313, D_F2_ = 2.683). However, this difference between anti-phase and in-phase modulation was abolished in hV4 (Wilcoxon signed-rank test hV4: p_F1 two-tailed_ = 0.469, D_F1_ = 0.816; p_F2 two-tailed_ = 0.125, D_F2_ = 0.849). The effects of attentional modulation on the anti-phase and in-phase oscillatory populations therefore differ most strongly in earlier compared to later stages of visual cortical processing.

The categorical definition of ROIs V1-hV4 may obscure a more continuous profile of attentional modulation on in-phase and anti-phase BOLD oscillations. Moreover, the magnitude of the BOLD response and the total number of vertices can differ between ROIs, which both complicate comparisons of modulatory effects between ROIs. To overcome these obstacles, we next examined the continuous relationship between attentional modulation and geodesic distance from V1 across participants, for the in-phase and anti-phase vertices separately (Fig. 4e,f), while also covarying for the amplitude of the BOLD signal during the unattended condition for each frequency (F1: uF1; F2: uF2) and proportion of anti-phase vertices within an ROI. The resulting Fisher-transformed spearman correlation is tested against 0 using a Wilcoxon signed-rank test in each phase group. We found a significantly positive monotonic relationship between attentional modulation and visual hierarchical distance in in-phase (p_F1 two-tailed_ = 0.016, D_F1_ = 1.936; p_F2 two-tailed_ = 0.016, D_F2_ = 1.657), but not anti-phase (p_F1 two-tailed_ = 0.934, D_F1_ = 0.022; p_F2 two-tailed_ = 0.813, D_F2_ = 0.388) populations. This supplements our observation that attention differentially modulates in-phase and anti-phase responses, and further shows that directed attention increasingly enhances the amplitude of in-phase, but not anti-phase, BOLD oscillations ascending the visual hierarchy.

### Attentional modulation of anti-phase BOLD oscillations predicts task performance

Altogether, our frequency-tagged BOLD correlates of stimulus enhancement and suppression, i.e. in-phase and anti-phase oscillations ^1^, respectively, exhibit dissociable profiles of attentional modulation which depend on the stage of visual cortical processing. The effects observed for in-phase and anti-phase oscillations are consistent with a more broadly distributed suppressive ‘field’ surrounding the hierarchical propagation of an enhanced signal ^12^.

We next asked whether attentional modulation of BOLD oscillations predicts task performance, and if so, how performance is shaped by the simultaneous enhancement (in-phase modulation) and suppression (anti-phase modulation) of competing visual cortical inputs. To do so, we first computed a metric of target detection sensitivity for each participant and for each run of the fMRI attention task, where a decrease in target contrast over trials of a run at a constant performance level of 80% indicates an increase in target sensitivity (Fig. 5a). For each participant, linear regression was then used to examine how, over runs of the task, the slopes of their target detection sensitivities over trials relates to their concurrent attentional modulation of either the in-phase or anti-phase BOLD oscillations. The within-participant correlation coefficients were then aggregated across individuals for random effects analyses. We found that over experimental runs of the task, larger improvements in target detection sensitivity were predicted by stronger attentional modulation of anti-phase BOLD oscillations across the visual cortex (Wilcoxon signed-rank test: p_two-tailed_ = 0.016, D = 1.765) but not in-phase BOLD oscillations (Wilcoxon signed-rank test: p_two-tailed_ = 0.156, D = 0.742; Fig 5b). In a direct comparison between in-phase and anti-phase populations, we found that the relationship of attention modulation with target sensitivity was significantly stronger for the anti-phase relative to in-phase populations (Wilcoxon signed-rank test: p_one-tailed_ = 0.039, D = 1.174).

**Figure 5.**
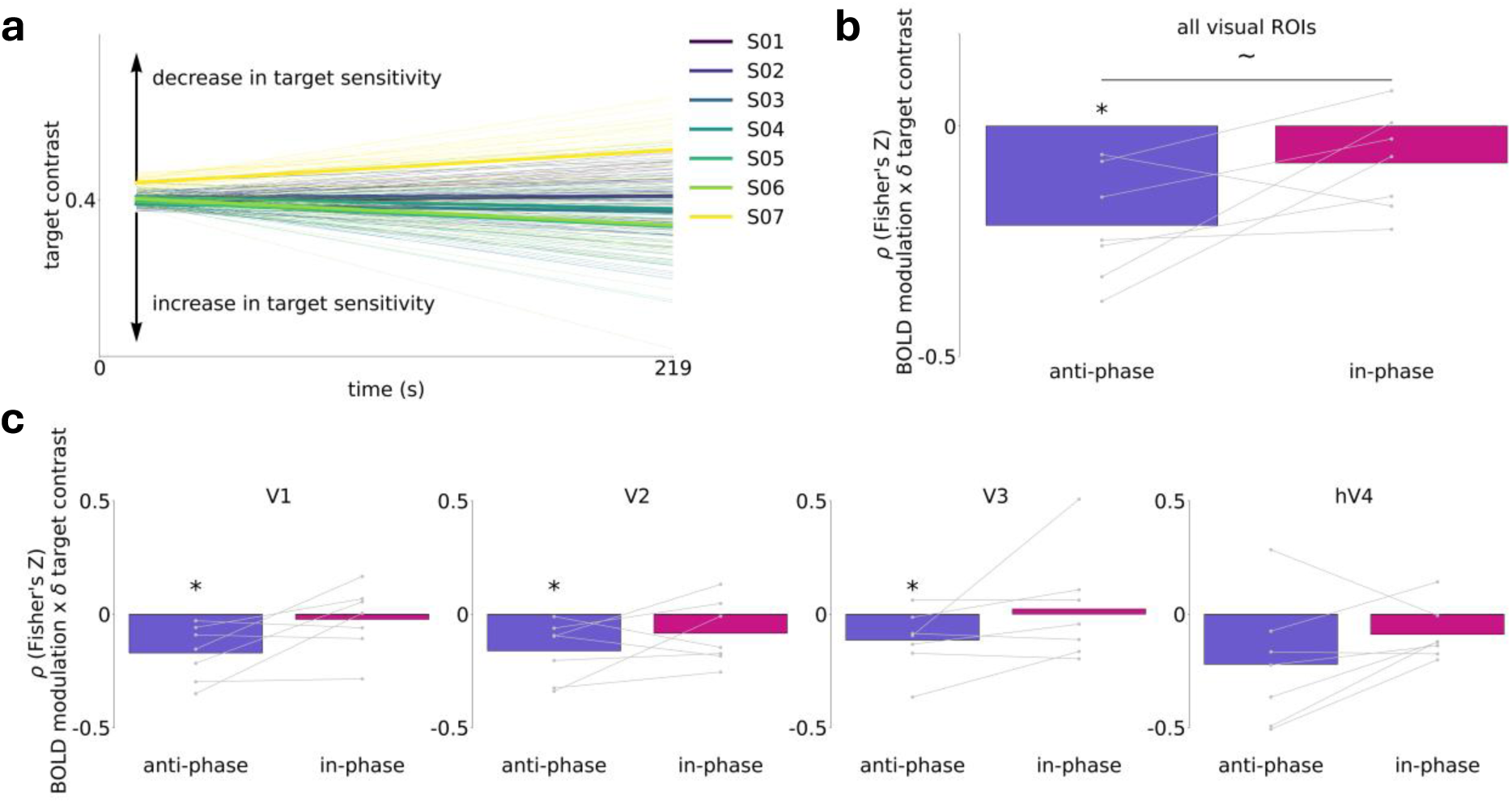
Attentional modulation of anti-phase BOLD oscillations predicts task performance. **(a)** Change in target sensitivity across experimental runs for each participant. For each experimental run, target sensitivity was assessed using the change in target contrast over the course of the run. Target sensitivity is summarized using the fitted linear slope of the changes in target contrast over the course of the run. Thick colored lines denote the mean slope across runs for each participant, and thin lines denote the estimated slopes for individual runs of each participant. **(b)** Run-by-run decreases in the fitted staircase slope (improvements in target sensitivity) relate to increases in the attentional modulation of anti-phase (slate-blue), but not in-phase (violet-red), oscillatory BOLD amplitudes. Gray dots represent the participant Fisher’s Z transformed spearman correlation values averaged across F1 and F2. **(c)** Relationship between target sensitivity and attentional modulation of oscillatory BOLD amplitude in individual visual ROIs. Representation of the data follows the same convention as in (b). Statistical significance was determined using a Wilcoxon signed-rank test (*P_two-tailed_ < 0.05; ∼P_one-tailed_ < 0.05).

Next we examined if the observed brain-behavior relationship between anti-phase modulation and target detection sensitivity is expressed uniformly across the visual cortex, or alternatively, exhibits a dependence on early versus later stages of processing. Consistent with the more distributed spatial profile of attentional modulation on anti-phase oscillations (Fig 4B), we observed that stronger attentional modulation of anti-phase responses predicted better target detection sensitivity across multiple visual cortical ROIs (Wilcoxon signed-rank test V1: p_two-tailed_ = 0.016, D = 1.401; V2: p_two-tailed_ = 0.016, D = 1.241; V3: p _two-tailed_ = 0.047, D = 0.849; hV4: p _two-tailed_ = 0.109, D = 0.802). This relationship was absent for in-phase vertices across all visual regions (Wilcoxon signed-rank test V1: p_two-tailed_ = 0.938, D = 0.157; V2: p_two-tailed_ = 0.156, D = 0.594; V3: p _two-tailed_ = 0.813, D = 0.006; hV4: p _two-tailed_ = 0.156, D = 0.749; Fig 5c).

## Discussion

In this study, we introduce the method of ft-fMRI to study the influence of directed attention on competing stimulus representations in the human visual cortex. We discovered that under sensory competition, stimulus-synchronized BOLD oscillations expressed throughout the visual cortex self-organized into spatially dissociated in-phase (enhancing) and anti-phase (suppressing) neuronal populations. The spatial topographies of these populations were highly replicable across scan sessions within participants, indicative of a fine-grained map of competitive feature-tuned responses.

Moving from striate to extrastriate visual cortex, directed attention increased the amplitudes of neuronal populations oscillating in-phase with the stimulus, consistent with prior event-related fMRI work ^10,11^. By contrast, the effects of directed attention on populations oscillating anti-phase with the stimulus were observed throughout the cortical hierarchy, consistent with a distributed suppressive field predicted by the normalization model and observed in electrophysiology work ^12–14^. The strength of this suppressive field, as indexed from individual differences in the amplitudes of anti-phase oscillations, significantly and selectively predicted psychophysical correlates of attentional performance.

The current study, along with our related ft-fMRI work ^2^ provide a major methodological advance for tracking competitive neuronal responses in humans. Frequency tagging stimulus evoked BOLD oscillations provides access to rich spatiotemporal dynamics including amplitude and phase information. We were able to demonstrate that the modulatory effects of directed attention on the amplitudes of frequency tagged BOLD oscillations depend both on their phase offset and location along the visual hierarchy. In a companion paper to the present work ^2^, we have also demonstrated that frequencies evoked by two competing unattended stimuli are detectable in the time series of individual voxels. These multiplexed oscillations synchronized to both stimulus frequencies at once, consistent with competitive interactions between different feature-tuned populations at subvoxel spatial resolution. Voxels were also detected which exhibited oscillations at intermodulation frequencies, which are produced from the nonlinear combinations of the fundamental frequencies of two stimuli, consistent with higher order integration of competing stimuli ^15–17^. Future work examining the effects of directed attention on multiplexing and intermodulation of BOLD oscillations in brain areas extending beyond visual cortex may shed light on the relationships of attention to cortical computation and communication at a fine-grained spatial scale.

Emerging electrocorticography evidence from pre-surgical epilepsy patients indicates attention operates via spatially and temporally dissociated mechanisms of alpha-band modulation ^18^. At visual field locations where participants focused their attention, the authors observed enhanced *cross-frequency coupling* between alpha-band (∼10 Hz) and higher-frequency bands along a feedforward pathway from V1 to dorsal intraparietal areas. By contrast, unattended visual field locations were associated with enhanced local intraareal coupling within ventrolateral areas but reduced interareal interactions along the visual hierarchy. Although alpha band oscillations are beyond the temporal upper bound of ft-fMRI (∼ 1 Hz) ^3,8^, our study provides strong complementary evidence for a network configuration whereby attention amplifies information flow for targets across the visual hierarchy while simultaneously limiting the transmission of neural signals across brain regions via a distributed suppressive field. Adding further evidence to this mechanistic dissociation, we show that individual differences in the strength of suppressive field modulation selectively predicted psychophysical performance on the attention task over and above in-phase (target) modulation (Supplementary Figure. 10). Altogether, these human electrocorticography and fMRI studies build on electrophysiology evidence from non-human primates suggesting that attentional enhancement and suppression of sensory information operate via distinct but complementary neuronal and biochemical bases ^14,19–25^.

The brain’s capacity to resolve competing sensory inputs via directed attention changes across its neurodevelopmental span, from infancy to older age ^26–28^, and in disorders which affect the neural and biochemical bases of attention such as Alzheimer’s disease, schizophrenia and autism ^29–31^.

For instance, both EEG and fMRI have revealed that compared to younger adults, older adults exhibit a reduced capacity to suppress unattended sensory information ^32–37^, leading to greater distractibility and unintentional encoding of information in environments characterized by a high degree of sensory competition. Based on our current findings in young adults, we speculate that under sensory competition in a ft-fMRI paradigm, older adults might exhibit a weakening of the suppressive field, indexed from altered spatiotemporal dynamics of anti-phase synchronized populations. The fine-grained spatiotemporal dynamics and extent of this cortical impairment remain to be determined, as they are invisible to standard event-related fMRI designs and EEG-derived SSVEPs. ft-fMRI provides a direct spatial and temporal isolation of the suppressive field for testing this and related predictions across neurodevelopmental stages and clinical conditions.

The current implementation of ft-fMRI has some limitations. In order to facilitate frequency tagging of BOLD oscillations up to 0.2 Hz, we used a fast acquisition sequence (TR=250 ms) which limited our field of view to a 16 slice slab centered on the visual cortex. Beyond our *a priori* ROIs in the striate and extrastriate visual cortex, we note that frequency-tagged BOLD oscillations were reliably detected in posterior parietal vertices at the edge of our field of view (Figure 1, Supplementary Fig. 11). These observations suggest the intriguing possibility that association and potentially even transmodal cortices might exhibit attention-dependent stimulus-synchronized BOLD oscillations. How the spatiotemporal dynamics of these higher-order cortical oscillations relate to the stimulus itself as well as to oscillations in the primary visual cortex remains to be determined.

In sum, we used ft-fMRI to provide direct in vivo evidence of competitive interactions among location-tuned neuronal populations in the visual cortex. We show that when two stimuli compete simultaneously for representational dominance, the phase offsets between oscillating stimuli and their frequency tagged cortical responses dissociate fine-grained maps of in-phase (enhancement) and anti-phase (suppression) BOLD oscillations. Modulation of these maps by directed attention is differentially distributed across the visual hierarchy and differentially related to psychophysical estimates of attentional performance. Looking ahead, ft-fMRI is a powerful tool for advancing a computationally informed understanding of the fine-grained spatiotemporal dynamics underlying human attention.

## Methods

### Participants

Eight right-handed participants (4 female) with normal or corrected-to-normal vision participated as paid volunteers after giving informed written consent. Their ages ranged from 20-31 years old (M = 26.25, SD = 3.49). Participants were compensated at a rate of CAD$15 per hour for their participation in the study. Participants also received an additional CAD$0.05 for each correct response during the behavioral task. One female participant was excluded from further analyses due to large gaze position deviations from fixation throughout the experiment (gaze area across experimental runs > 5 degrees^2^). All experimental procedures were approved by the Health Sciences Research Ethics Board at Western University.

### Stimuli & Task

We generated the experimental stimuli and tasks using PsychoPy ^38^. Stimuli were projected at 60 Hz onto a screen (resolution of 1024x768 pixels) fixed at the back of the scanner bore from a frame rate synchronized stimulus laptop. Participants viewed the projected visual stimuli via a mirror fitted into the head coil placed in front of the eyes (total distance to screen is 84 cm).

We presented participants with a pair of wedge stimuli occupying one visual field quadrant and another pair of radial wedge stimuli occupying equal spatial portions of the opposing visual field quadrant. Each wedge was part of a checkerboard pattern. The wedge stimuli extended from 1.5° to 10° eccentricity. Participants initiated each experimental run via a button press. During a 14s baseline period at the beginning of each experimental run, we used a red line(s) to cue our participants to covertly attend to the wedge(s) in one visual field quadrant. Following cue offset, participants had to covertly attend to and detect transient color changes (lasting for 500 ms) at either (directed attention task) or both (distributed attention task) oscillating wedge stimuli presented in either the upper right or upper left visual field quadrant. Oscillating wedge stimuli were sine-squared modulated at 0.125 Hz and 0.2 Hz, and exhibited a 12 Hz counterphase flicker.

Color changes only occurred when the oscillating stimuli were at above 33% luminance contrast. Color change inter-stimulus-interval was adjusted for the frequency of the oscillating wedges. For the stimulus oscillating at 0.125 Hz, color changes occurred at 0.5-2 s pseudo-random intervals drawn from a uniform distribution. For the stimulus oscillating at 0.2 Hz, color changes occurred at 0.5-1.25 s pseudo-random intervals drawn from a uniform distribution. Color changes during the directed attention conditions had a 50% chance of occurring on each oscillation of the wedge. The participant had 750 ms to respond to the color change occurrence via button press with the right index finger. If the response did not occur within the response interval, the response was recorded as a false alarm. If the participant did not respond at all before the next color change appeared, this was recorded as a missed trial. Color change intensity was continuously adjusted on each experimental run using an adaptive staircase procedure (QUEST) ^39^ to achieve a response accuracy of approximately 80%, where response accuracy was calculated as the number of correct responses divided by the total sum of correct responses, false alarms and missed trials. QUEST parameters: ß (slope of psychometric function) = 3.5; δ (probability of incorrect response under ideal conditions) = 0.01; γ (probability of false alarm) = 0.05; Reference stimulus color change = 40% of maximum contrast.

Each participant performed a total of sixteen 219s runs of each of the three experimental conditions; ‘distributed-attention’, ‘attend-F1’, ‘attend-F2’, over the course of four experimental sessions (4 runs of each condition per session). We counterbalanced the location of the stimuli in either the upper right or upper left visual hemifield across participants (4 participants per location).

### MRI acquisition

MRI data were acquired on a Siemens Magnetom 7T machine (MAGNETOM Prisma, Siemens Healthineers, Erlangen, Germany) at the Center for Functional and Metabolic Mapping using a Siemens 32-channel whole-head array for the anatomical T1-weighted images and a custom occipital-parietal 32-channel head coil for the functional T2/T2*-weighted images.

A T1-weighted anatomical MP2RAGE scan was acquired during the first study visit for registration purposes and to enable cortical surface projection (TR 6000 ms, TE 2.74 ms, TI1 800 ms, TI2 2700 ms, FOV 240 mm, flip angle 1 4°, flip angle 2 5°, 0.7 mm isotropic voxels, 224 slices, AP phase encoding direction, in-plane acceleration factor 3, slice partial Fourier 6/8). fMRI scans were collected during each of the four study visits using a multiband T2*-weighted echo-planar imaging sequence covering an oblique slab centered on the ventral occipital lobe (TR 250 ms, TE 20 ms, FOV 208 mm, flip angle 30°, 2.5 mm isotropic voxels, 16 slices, RL phase encoding direction, no in-plane acceleration, and multi-band acceleration factor 4) with 880 volumes acquired per scan for a total acquisition time of 220s. T2-weighted anatomical images (TR 2,000 ms, TE 20 ms, FOV 256 mm, flip angle 30°, 2 mm isotropic voxels, 80 slices, RL phase encoding direction, in-plane acceleration factor 2, and multi-band acceleration factor 4) were acquired in the middle of each study session to facilitate the registration between session-specific slab fMRI data and the T1-weighted anatomical image.

Data were converted from dicom to nifti brain imaging data structure (BIDS) format using heudiconv (v0.11.3) ^40^ and dicom2niix ^41^.

### MRI data processing

MRI data was preprocessed using a pipeline developed with *Nipype*, which used the following software packages: ANTS, Freesurfer, FSL, AFNI, workbench, and customized Nipype workflows adapted from smriprep and fmriprep ^2^. Existing Nipype workflows from fmriprep were customized to integrate session-specific whole-brain EPIs and run-specific EPIs into the registration workflow.

To preprocess each fMRI dataset, slice-timing correction, followed by single-shot interpolation to T1w space, surface resampling to 32k vertex fsLR surfaces (fsLR 32k surfaces), and nuisance regression were performed. fMRI data was regressed using 24 motion parameters (six HMC parameters and their 1st and 2nd order first derivatives), high pass filter regressors (<.01 Hz), and the mean WM and CSF signal to minimize the effects of scanner drift, motion and other non-neural physiological noises. Each fMRI run was truncated between 59 and 219 seconds (or 45 seconds after the oscillatory stimulus begins and when the stimulus ends) to account for transient hemodynamic effects associated with the onset of continuous oscillatory stimulation, and to ensure that the run duration provided the frequency resolution necessary for precise phase estimation at the frequency of both oscillating wedge stimuli. Since our stimuli had periods of 8s (0.125 Hz) and 5s (0.2 Hz), we were able to obtain 20 and 32 full cycles, respectively, at our frequencies of interest from the resulting pre-processed and noise-corrected 160s time course.

Details pertaining to resampling of individual fMRI runs to fsLR 32k surfaces are as follows: The participant-specific T1-weighted (T1w) structural image was skull-stripped using SynthStrip and then processed with smriprep to segment the grey matter (WM), white matter (GM), and cerebrospinal fluid (CSF). Subsequently, the T1w image was registered to the single-band reference (SBRef) image from each scan-specific whole-brain EPI using a boundary-based registration (BBR) cost function. The cortical ribbon mask was transformed into the whole-brain EPI space for each session and utilized to perform a BBR-based registration between the SBRef images of the scan-specific whole-brain EPIs and their corresponding scan-and-run-specific slab EPIs. N4 bias field corrections were performed to each image prior to performing the registrations. Next, after applying low-pass filtering to each fMRI dataset, head motion correction (HMC) was performed using FSL’s MCFLIRT, registering all volumes to their associated run-specific SBRef image. We low-pass filtered the HMC parameters to remove unwanted fluctuations (0.2-0.4 Hz) in the estimated HMC time courses. Note that low-pass filtered data was used solely to generate fluctuation-free HMC transformations, which were then applied to the raw, unfiltered fMRI data. To enable one-shot interpolation of each fMRI run to T1w space, the transformations were concatenated in the following order: HMC to run-specific slab EPI, slab EPI to session-specific whole brain EPI, and whole brain EPI to participant-specific T1w image. Next, the volumetric BOLD data (in T1w space) was resampled onto the participant’s native surface, and subsequently resampled onto a standard fsLR 32k surface.

### Activation thresholding

Preprocessed fMRI runs were grouped for each participant for the localizer condition (distributed-attention task) to generate frequency-encoded brain maps. A Monte Carlo random subsampling approach was used to ensure that the localized frequency-encoded populations were robust against random subsamples. This involved performing 500 iterations of model fitting. In each iteration, preprocessed fMRI runs from a single participant were randomly split into train and test groups (50/50 split). An average fMRI run was computed for each data split, followed by time series normalization to BOLD % change relative to the mean of the time series, and subsequent fitting with a set of frequencies.

To generate a frequency-encoded map, a general linear model was fitted on each iteration using a Fourier basis set at each frequency (F1 = 0.125 Hz; F2 = 0.2 Hz; see *Fitting frequency-encoded populations*). Maps of each frequency were binarized by thresholding with an unadjusted p-value < 0.05 (F-test). These activation maps were aggregated across all 500 iterations by computing the mean, producing a fractional overlap map that indicates the stability of frequency-encoded brain maps across random subsampled iterations. A value of zero means the frequency was not encoded in any of the 500 random subsamples, while a value of one means it was encoded in all 500 subsamples. Based on our previous work ^2^, we thresholded frequency-encoded maps at a fractional overlap value of 0.8 (i.e. vertices exhibiting significant frequency-specific modulation in at least 400/500 subsamples).

### ROI definition

For each participant, we selected ROIs from a visual surface-based cortical parcellation ^7^. We specifically selected ROIs that were targeted by our stimulation protocol (i.e. those located on the ventral visual surface: V1v, V2v, V3v, hV4). We only included cortical hemisphere-specific data from an ROI in our analysis if we could on average localize at least 25 suprathreshold vertices (fractional overlap > 0.8) across resamplings of the data in that hemisphere.

### Fitting frequency-encoded populations

To model a frequency of interest in the fMRI data, vertex-wise BOLD time series were fitted using a general linear model. The model included sine and cosine regressors associated with each frequency of interest, and an F-test was performed on the combined fit to the sine and cosine regressors at each frequency of interest to test for a frequency-specific response. The amplitude and phase of the frequency-specific response was then determined using the coefficient weights of the sine and cosine components at each frequency. Description of the general linear model is formally described below.

For any given BOLD time series, the general linear model can be expressed as follows:

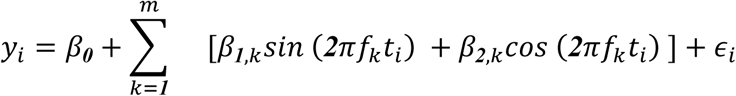

where:

- *β_0_* is the intercept term
- *β_1_*_,*k*_ and *β_2_*_,*k*_ are the coefficients associated to the sine and cosine terms of the k^th^ frequency *f*_*k*_, respectively
- *f*_*k*_ is the k^th^ frequency of interest
- *ϵ*_*i*_ is the error term for observation *i*
- *m* is the number of frequencies of interest

The amplitude (*BOLD*_%*k*_) at a frequency of interest was calculated as the root sum squared of the fitted coefficients associated with the Fourier basis of that frequency. This value was then multiplied by 2 to include the negative portion of the oscillatory response in the amplitude (peak-to-trough) estimate. For each participant, task condition, and *k^th^* frequency of interest, we obtained 500 estimates of the response amplitude via the iterative resampling procedure described in *Activation thresholding*. The reported amplitude estimate for each vertex corresponds to the mean amplitude across resamplings of the data.

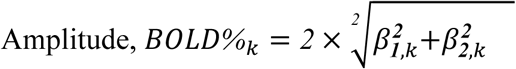

The phase-offset (*θ*_*k*_) of a frequency of interest was calculated as the arctangent of the Fourier basis coefficients of that frequency. For each participant, task condition, and *k^th^* frequency of interest, we obtained 500 estimates of the phase offset via the iterative resampling procedure described in *Activation thresholding*. The reported phase estimate for each vertex corresponds to the circular mean of the phase estimates across resamplings of the data.

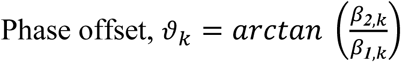

### Classification of in-phase and anti-phase synchronized responses

To separate vertex-wise data into populations that synchronize in-phase and anti-phase with the stimulus, we fit a von Mises mixture model to the phase offset data estimated from the localizer (distributed-attention task) using the *clustering* algorithm from the *pycircstat2* library. This resulted in two unique phase distributions for each frequency of interest. To determine which of these distributions described in-phase or anti-phase responses, we first convolved the stimulus time course at each frequency with a gamma hemodynamic response function (HRF), and truncated the convolved time course to the period of the analysed data (that is, 59s - 219s). Next, we calculated the phase of the resulting time course to obtain an estimate of the in-phase response predicted by the canonical HRF. We computed the circular distance between each distribution and the predicted HRF phase by obtaining the average circular distance across vertex phase estimates within a distribution and the predicted HRF phase. We then defined vertices falling within the phase distribution with the smallest circular distance to the predicted HRF phase as responding in-phase with the stimulus. Vertices falling within the opposing distribution were defined as responding out of phase (anti-phase) with the stimulus.

### Attentional modulation analysis

To obtain an estimate of attentional modulation at our frequencies of interest, we first calculated the difference in BOLD amplitude between the condition in which the frequency of interest was attended relative to when the frequency was unattended across 500 resamplings of the data in each of these conditions (F1 attentional modulation = aF1_BOLD%_ - uF1_BOLD%_; F2 attentional modulation = aF2_BOLD%_ - uF2_BOLD%_). We then averaged the attentional modulation values across estimates to obtain the reported measure of attentional BOLD amplitude modulation.

To test for significant attention-driven changes in BOLD amplitude, we contrasted maps of BOLD amplitude between the condition in which the frequency of interest was attended versus the condition in which the frequency was unattended at each frequency of interest (aF1_BOLD%_ versus uF1_BOLD%_; aF2_BOLD%_ versus uF2_BOLD%_). For each vertex, we performed the contrast using a two-tailed t-test between the aggregated run BOLD amplitude data corresponding to each of the compared conditions. Contrasts were performed again after shuffling the condition labels between each of the contrasted conditions. Condition labels were re-shuffled 1,000 times for each contrast to generate a null distribution of t-statistic values for each vertex. The order of the vertex-wise t-value of the original contrast was then compared against this null distribution, and t-statistic maps were thresholded for vertices exhibiting t-statistic values at p < 0.05.

### Attentional modulation as a function of geodesic distance from V1

Surface geodesic distance from V1 was used to obtain estimates of visual hierarchical distance in localized vertices. We first used the workbench (v1.5.0) *wb_command -surface-geodesic-distance-all-to-all* method to create a map of geodesic distances between all vertices on the surface of each cortical hemisphere. We anatomically constrained this map to V1v as defined by a visual surface based cortical parcellation ^7^, and computed the mean geodesic distance between V1v vertices and the rest of the cortical surface. Next, we examined attentional modulation at F1 (aF1_BOLD%_ - uF1_BOLD%_) and F2 (aF2_BOLD%_ - uF2_BOLD%_) as a function of geodesic distance in V1-hV4 for suprathreshold in-phase and anti-phase vertices separately. The partial correlation between attentional modulation and geodesic distance was then computed using *partial_corr* from the *pingouin* library, where the proportion of anti-phase vertices within an ROI, and the amplitude of the the BOLD signal during the unattended condition, were used as covariates to isolate the unique relationship between attentional modulation and V1 geodesic distance.

### Assessing participant phase, signed phase, and attentional modulation map reliability

For each participant, we acquired data for each of our task conditions over the course of 4 separate experimental sessions. To test for the reliability of the reported task metrics at each frequency, we inspected the consistency of phase offset and signed phase maps under our localizer condition (distributed-attention task) condition, and the consistency of attentional modulation maps under our directed attention (Attend-F1: aF1-uF1 and Attend-F2: aF2-uF2) conditions, across experimental sessions. We first generated session specific phase and attentional modulation maps estimated from the run-average data of the distributed-attention and directed attention tasks, respectively, for each experimental session. These maps were thresholded according to the criteria described in *Activation thresholding* (i.e. fractional overlap > 0.8). We then examined the correspondence (circular correlation for phase maps, Dice’s coefficient for signed phase maps, and Pearson’s *r* for attentional modulation maps) between thresholded session-specific maps within the anatomically constrained visual ROI (*ROI definition*). To test for the significance of map correspondence across experimental sessions, we employed a generative modeling procedure since surface maps of task metrics are spatially autocorrelated (*BrainSMASH*) ^42^. For each participant, we used each session’s thresholded metric maps to generate 50 simulated maps that retain the spatial autocorrelation structure of the original map, resulting in a total of 200 surrogate maps (50 for each of the 4 sessions). The correspondence between each session’s map with these 200 surrogate maps within the anatomically constrained ROI was then computed to generate a distribution of 800 surrogate correlation (circular or Pearson’s) or overlap (Dice’s coefficient) values (4 session maps x 200 surrogate maps). We tested for the reliability of participant metric maps by comparing the distribution of between-session map correspondence values to the surrogate distribution. The reported test statistic was obtained by aggregating the within-participant between-session distributions of map correspondence values and the corresponding surrogate distributions across participants before performing the Kolmogorov-Smirnov (K-S) test to test for differences between these aggregated distributions, and the Mann-Whitney (M-W) U test to test for statistical differences between the distribution means.

To examine whether the observed spatial topographies of our metrics are specific to each participant, we compared within-participant between-session map correspondence to between-participant session map correspondence. To generate the between-participant distribution, we computed the correspondence (circular correlation for phase maps, Dice’s coefficient for signed phase maps, and Pearson’s *r* for attentional modulation maps) between within-participant session maps and session maps for participants who were presented with the same stimulus configuration (i.e. those presented with the identical positioning of oscillating wedge stimuli in the same visual field quadrant), thresholded using the same anatomical mask applied to the within-participant session maps. We then compared the distribution of within-participant session map correspondence values against the distribution of between-participant session map correspondence values. The reported test statistics were obtained by aggregating the within-participant distributions and between-participant distributions and performing the K-S test for differences between these distributions, and performing the M-W U test to test for statistical differences between the distribution means

### Eye-tracking data acquisition & analysis

Eye tracking was performed using an Eyelink 1000 system during the experiment. Eye tracking data was sampled at 0.5 kHz. Before each experimental session, eye position was calibrated using a 9-point calibration. For each experimental run, we obtained a time series of horizontal and vertical eye positions. Blinks were automatically detected by the Eyelink system and manually selected based on outlier values, and data within ±0.1 seconds of each blink were excised. We detrended recorded eye-positions to account for instrumental drift over the course of the experimental session. We plotted a 2D histogram of horizontal and vertical eye-positions for every experimental run. Eye positions were summarized by fitting a 2D Gaussian probability distribution to the data and calculating the area of the contour that contained 95% of the fitted distribution, and gaze area for each condition was computed as the mean area of the fitted distribution across runs.

### Behavioral data acquisition & analysis

During each experimental run, we recorded participant button press responses to a target detection task (see *Stimuli & Task*). Participant behavioral performance on the task was determined as the number of correct responses divided by the total sum of correct responses, false alarms and missed trials.

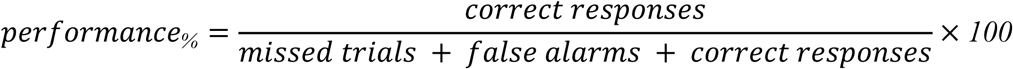

Behavioral performance during the experiment was controlled at 80% using an adaptive staircase procedure to ensure that participants were engaged at the attended location. The adaptive staircase registers improvements in target sensitivity as decreases in target contrast and vice versa. Hence, in each experimental run, we assessed improvements in target sensitivity by monitoring changes in the target contrast over the course of the experimental run. We fitted a linear regression model to target contrast changes over the course of an experimental run to summarize changes in target sensitivity using the slope of the model fit. A positive slope would indicate a decrease in target sensitivity over the course of a run (i.e. an increase in target contrast) and a negative slope would indicate an increase in target sensitivity over the course of a run (i.e. a decrease in target contrast). Using this information, we examined if there exists a relationship between target sensitivity and attention-driven changes in the amplitude of in-phase and anti-phase BOLD oscillations. For each participant, we computed the run-by-run correlation (Spearman’s ⍴) between the fitted target contrast slope (metric representing the inverse of target sensitivity) and BOLD amplitude values for the corresponding directed attention task separately at each frequency (i.e. Attend-F1 for F1 amplitude and Attend-F2 for F2 amplitude). We then averaged the Fisher’s Z transformed correlation values obtained at F1 and F2 for each participant before performing a random-effects analysis to determine if the relationship between target sensitivity and attentional modulation of the BOLD amplitude is significantly different from zero. This analysis was repeated after including the amplitude of the opposing phase group (i.e. run-wise amplitude of in-phase BOLD oscillations was included as a covariate when computing the partial run-wise correlation between the amplitude of anti-phase BOLD oscillations and psychophysical performance, and vice versa) as a covariate to ensure that the highlighted effects were specific to the examined phase group.

### Statistical analysis

Statistical analysis was performed using the *scipy* and *statsmodels* libraries in python. For vertex-wise inference, the *statsmodels* library was used to fit the GLM to the time series data and perform the F-test to determine the significance of frequency-specific responses. For participant-wise statistical inference, no assumptions were made about the normality of compared data distributions. The *scipy* library was used to perform the reported non-parametric statistical tests. An alpha level of p < 0.05 was used to assess significance.

## Data & Code Availability Statement

Example data and analysis code will be made available upon the publication of this manuscript, or via reasonable request to the corresponding author.

## Notes

### Competing Interest Statement

The authors have declared no competing interest.

