## Supplementary Materials for "Attentional enhancement and suppression of stimulus-synchronized BOLD oscillations"

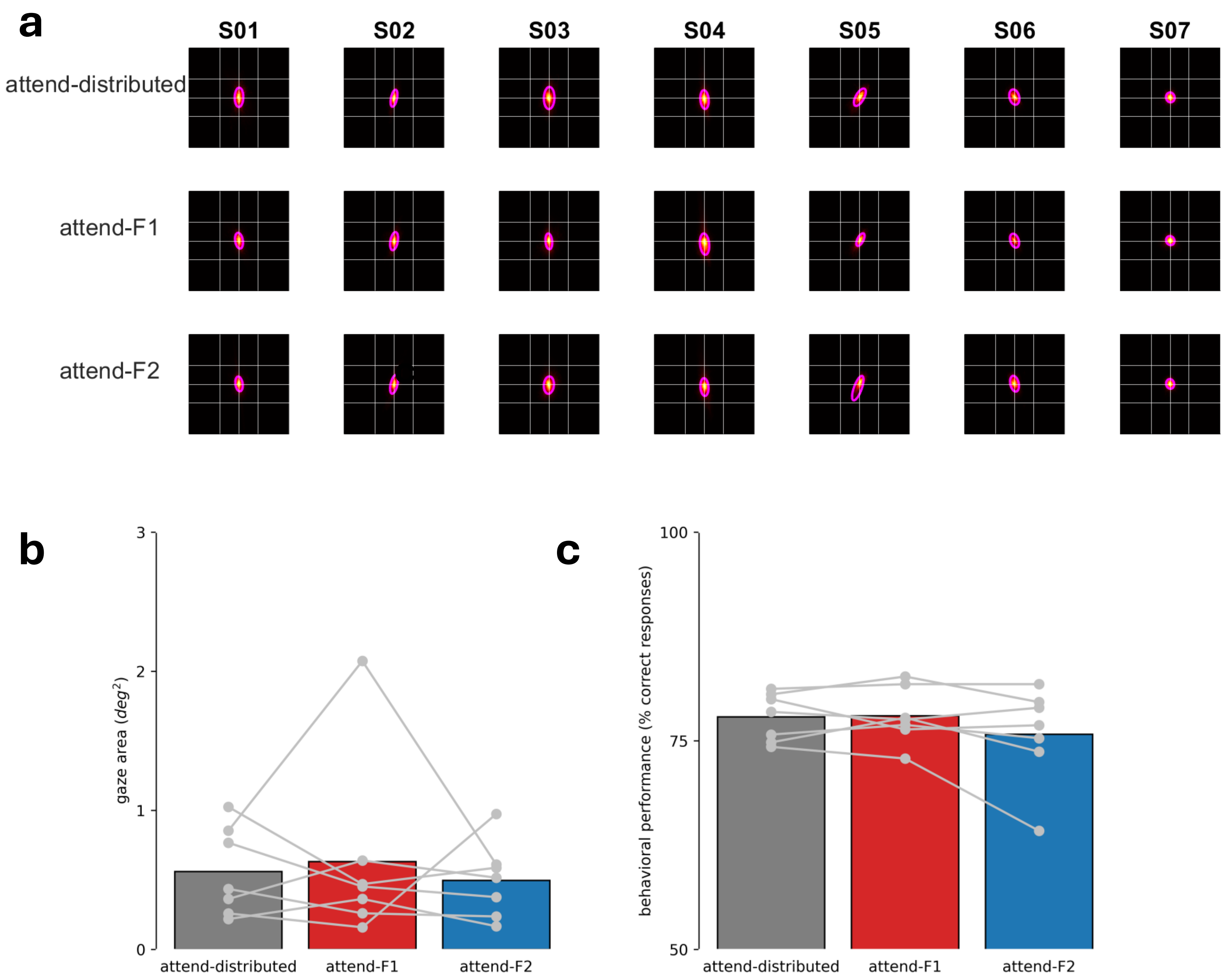


***Supplementary Figure 1. Participant behavioral performance across experimental conditions***

***(a)*** *Eye positions remain consistently near the center of the display. Eye-tracking data is summarized by displaying the heatmaps and fitted gaussian distribution of the aggregated data for each participant (columns) and experimental condition (rows). The grid lines indicate the radius of the central fixation region. The magenta contour lines indicate the area of the gaussian mixture model that contains 95% of the fitted data.* ***(b)*** *The gaze contour areas are comparable between experimental conditions across participants (K-W ANOVA; H(2) = 0.052; $p_{two\text{-}tailed}$=0.974). Data points indicate the participant mean gaze contour area across runs for an experimental condition. Data points are within the area of the central fixation region (~1.8 $deg^2$). Individual participants showed comparable gaze-areas between experimental conditions across runs (one-way ANOVA: $p_{two\text{-}tailed}$ > 0.05 for 7/7 participants).* ***(c)*** *Participant behavioral performance was comparable between experimental conditions across participants (K-W ANOVA; H(2) = 0.586; $p_{two\text{-}tailed}$ = 0.746). Data points indicate the participant mean behavioral performance across runs for an experimental condition. These data show that the adaptive staircase procedure maintained behavioral performance at ~80%. Individual participants showed comparable behavioral performance between experimental conditions across runs (one-way ANOVA: $p_{two\text{-}tailed}$ > 0.05 for 6/7 participants).*

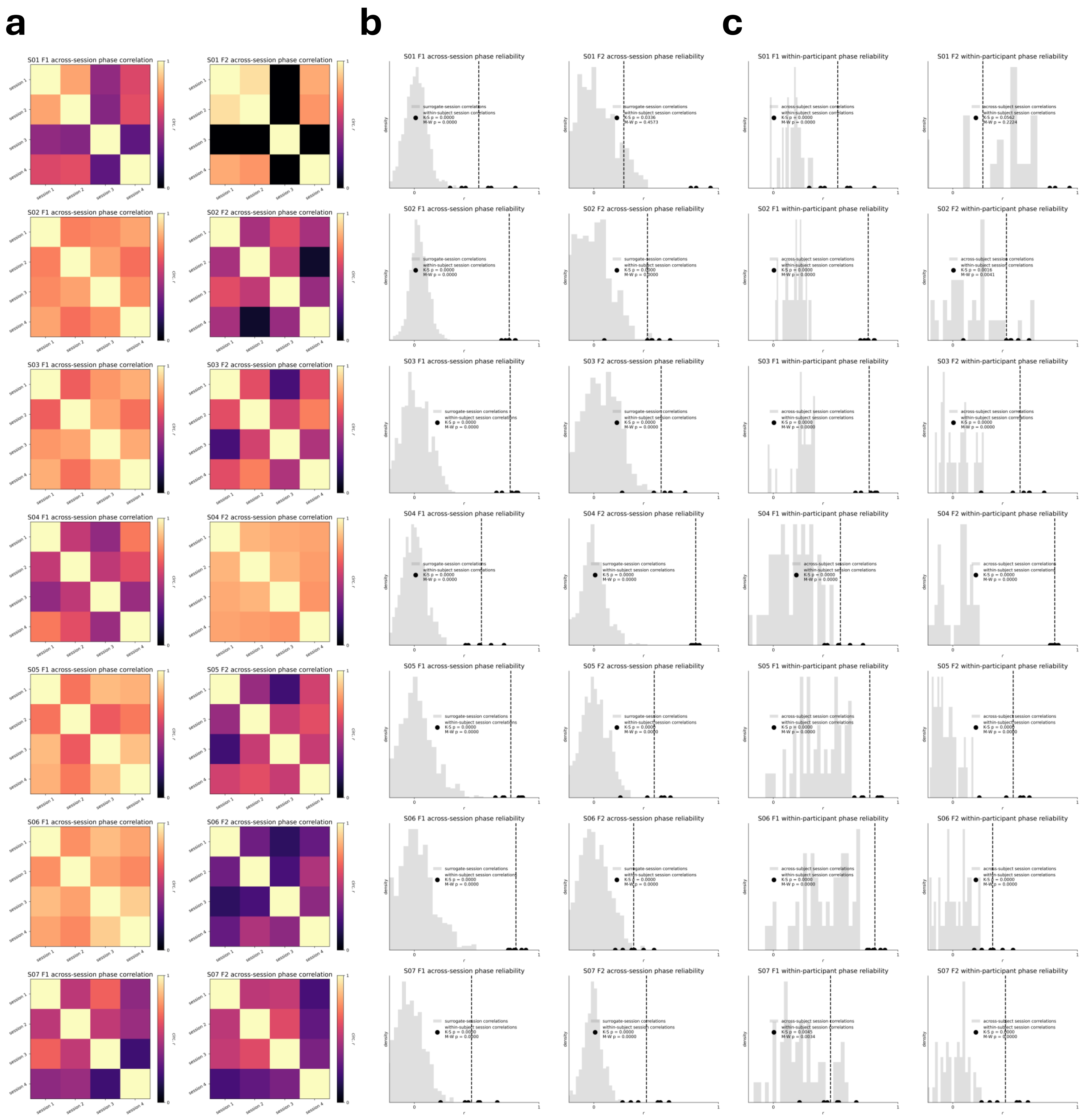


***Supplementary Figure 2. Reliability of phase maps across sessions and within participants***

*__(a)__ Correlation (circular r) between phase maps estimated from each experimental session (4 sessions; 4 runs of data each) for each participant (rows) at F1 (left) and F2 (right). __(b)__ Surrogate map test to determine the significance of between session correlations. Data in gray shows the distribution of surrogate correlation values generated by correlating session phase maps with shuffled phase maps that retained the same spatial autocorrelation structure and anatomical distribution as the original F1 (left) and F2 (right) session phase maps. Black data points represent the between session phase map correlations within participants. Dashed vertical lines indicate the mean value of between session phase map correlations. Reliability of phase maps was tested by comparing the between session correlations against the surrogate distribution for each participant (Kolmogorov-Smirnov $p_{one\text{-}tailed} < 0.05$ for 7/7 participants; Mann-Whitney U $p_{one\text{-}tailed} < 0.05$ for 7/7 participants at F1, and 6/7 participants at F2). This test was repeated for the between session correlations and surrogate distributions aggregated across all participants (Kolmogorov-Smirnov $p_{F1\ one\text{-}tailed} = 9.65 \times 10^{-60}$; $p_{F2\ one\text{-}tailed} = 2.45 \times 10^{-31}$; Mann-Whitney U $p_{F1\ one\text{-}tailed} = 3.66 \times 10^{-29}$; $p_{F2\ one\text{-}tailed} = 6.31 \times 10^{-21}$). __(c)__ Between participant session phase map test to examine the uniqueness of the functional topography of within-participant session phase maps. Data in gray shows the distribution of between-participant session phase map correlation values. Data representation follows the same convention as in (b). Individuality of phase maps was tested by comparing the within-participant between session correlations against the between-participant between-session correlations (Kolmogorov-Smirnov $p_{one\text{-}tailed} < 0.05$ for 7/7 participants; Mann-Whitney U $p_{one\text{-}tailed} < 0.05$ for 7/7 participants at F1, and 6/7 participants at F2). This test was repeated for the within-participant between session correlations and between-participant between session correlations aggregated across all participants (Kolmogorov-Smirnov $p_{F1\ one\text{-}tailed} = 1.03 \times 10^{-20}$; $p_{F2\ one\text{-}tailed} = 5.60 \times 10^{-23}$; Mann-Whitney U $p_{F1\ one\text{-}tailed} = 5.46 \times 10^{-20}$; $p_{F2\ one\text{-}tailed} = 1.49 \times 10^{-16}$).*

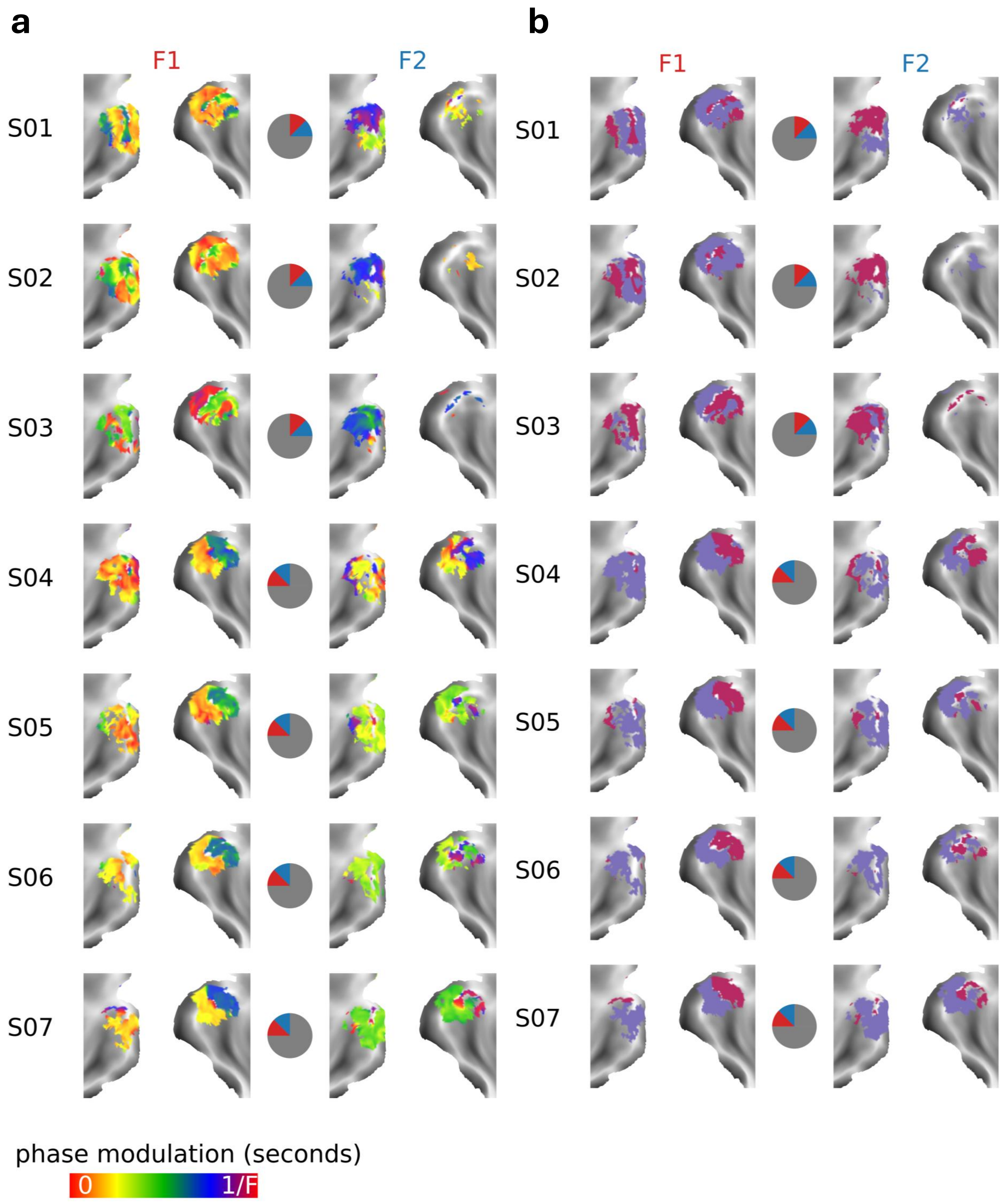


***Supplementary Figure 3: Individual participant phase maps and signed phase maps***

***(a)*** *Participant phase maps at F1 (left) and F2 (right). The color bar denotes the response phase scaled to the period of the examined frequency. The central diagram indicates the location of the oscillating wedge stimuli in the stimulated visual field quadrant for a given participant (F1 in red; F2 in blue).* ***(b)*** *Participant phase maps classified into in-phase (violet-red) and anti-phase (slate-blue) synchronized populations. Data arranged according to the same conventions as part (a).*

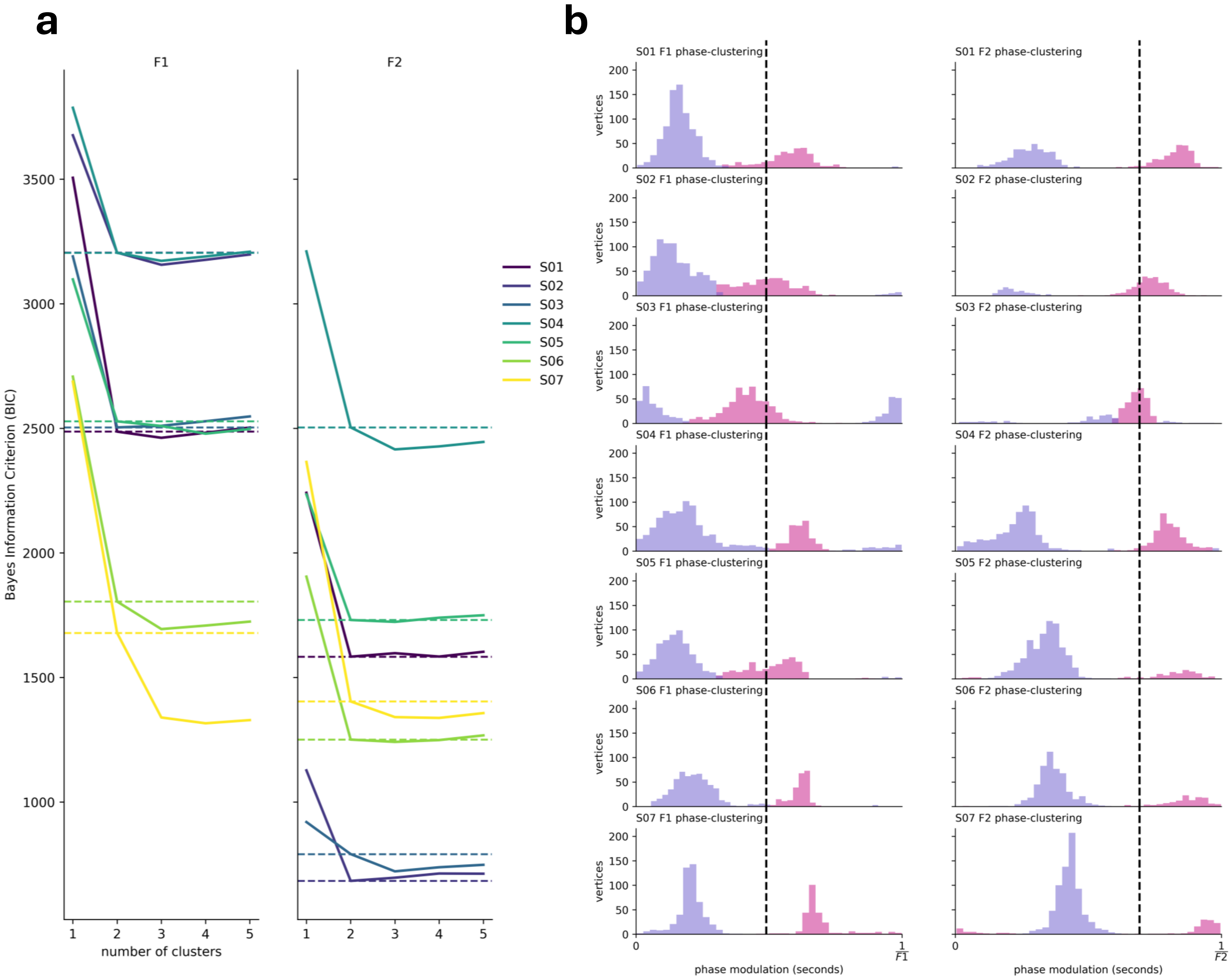


***Supplementary Figure 4: von Mises mixture model Bayes Information Criterion with increasing number of clusters, and individual participant phase clustered distributions***

***(a)*** *Individual participant data showing changes in von Mises mixture model information with an increasing number of phase clusters. Model information was assessed using the Bayes Information Criterion (BIC), where decreases in BIC indicate an increase in the explanatory power of the model. We fit a von Mises mixture model with an increasing number of clusters to the localizer phase data and found that 2 clusters maximize the information that the model can explain. The dashed horizontal line indicates the BIC value at 2 phase clusters. Changes in model BIC are marginal beyond 2 phase clusters.* ***(b)*** *Individual participant data (rows) showing the clustering of vertex phase responses with respect to the phase predicted by the HRF (black vertical dashed line) into in-phase (violet-red) and anti-phase (slate-blue) populations at F1 (left) and F2 (right).*

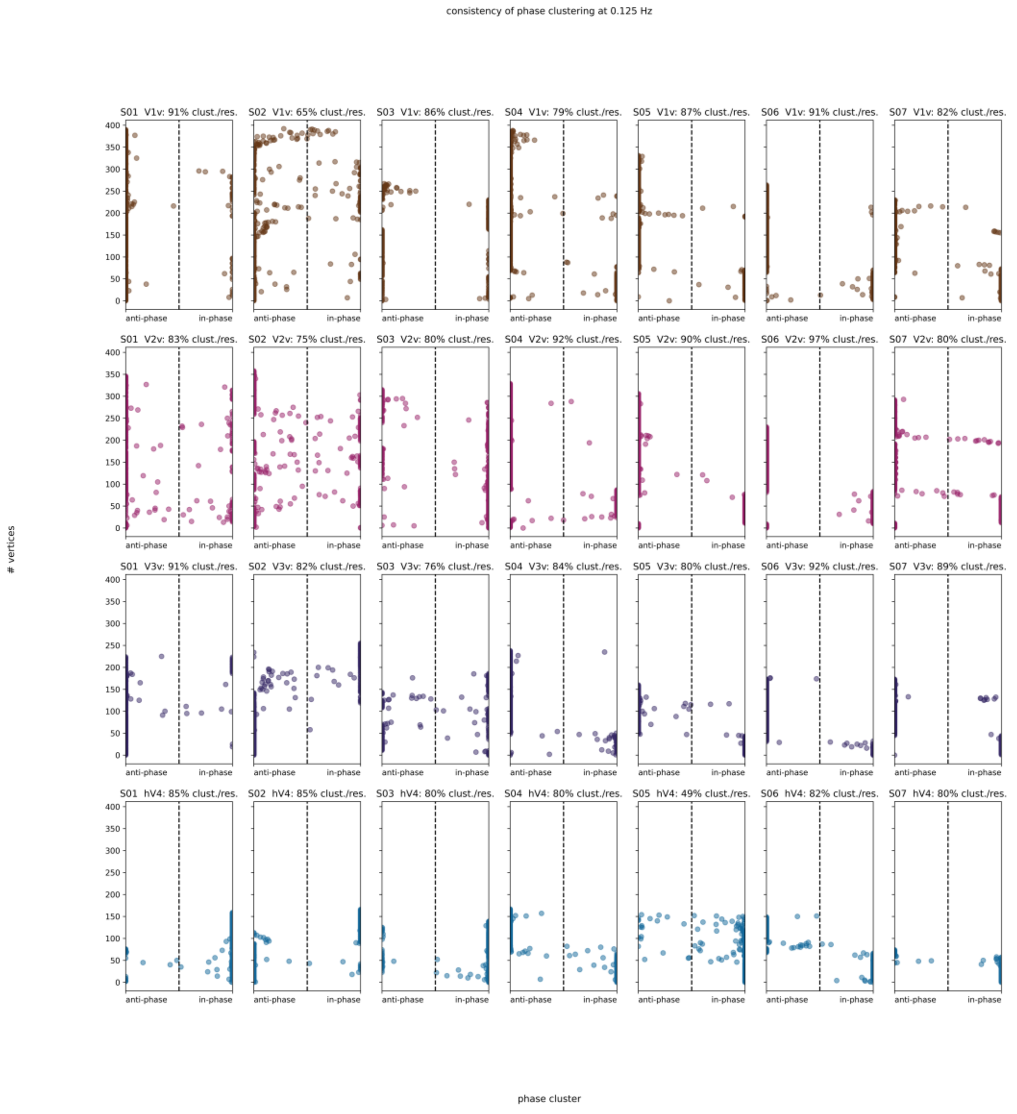


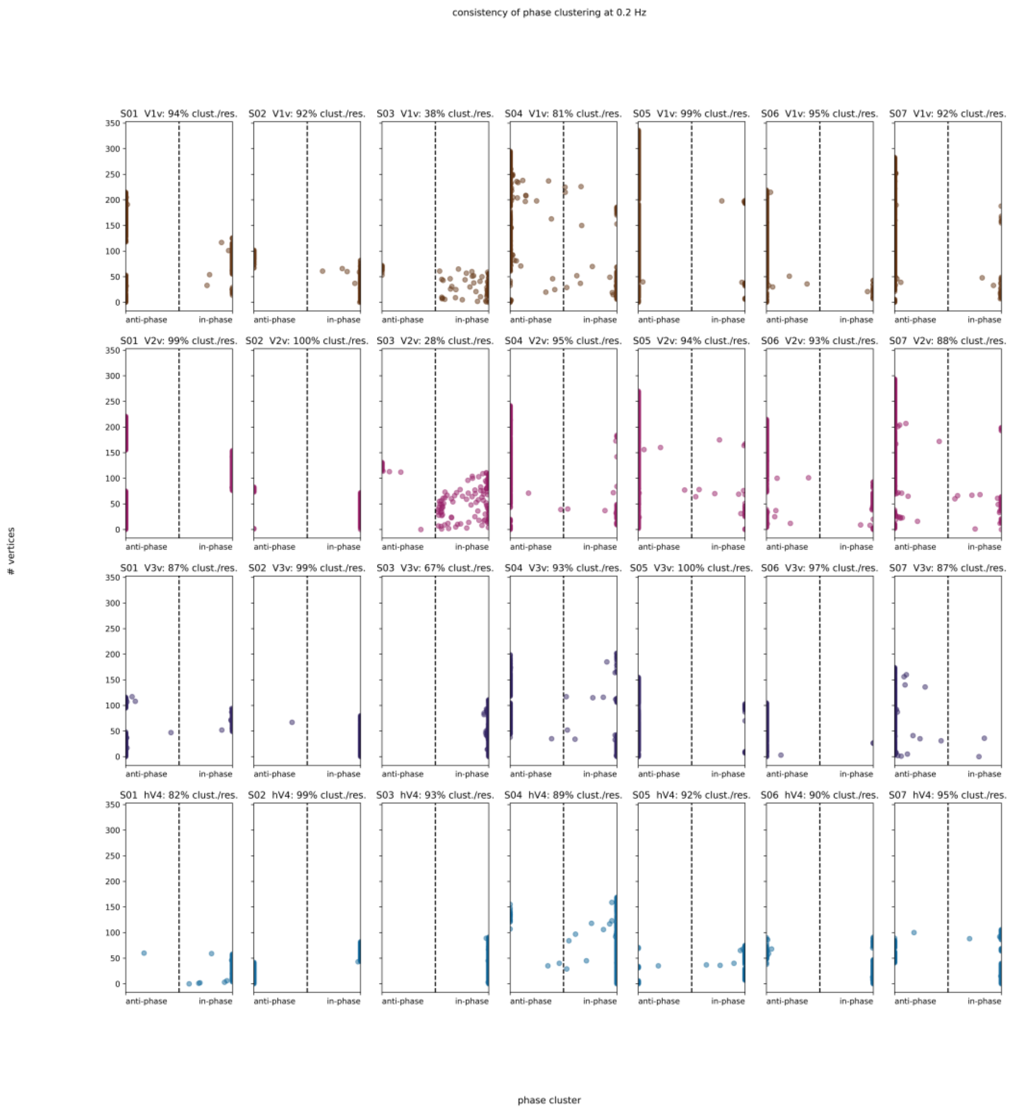


***Supplementary Figure 5: Reliability of phase clustering across random resamplings of the data***

*Individual participant data showing phase clustering reliability across participants (columns) and ROIs (rows) at F1 (top) and F2 (bottom). We examined the phase cluster assignment for each vertex across resamplings of the data (500 repetitions), and we show the proportion of bootstraps in which a vertex was assigned to one of the two phase clusters (in-phase or anti-phase). Vertices falling closer to the edges indicate a consistent phase cluster assignment across bootstraps; whereas vertices falling closer to the middle (dashed vertical line) indicate low reliability in phase cluster assignment across bootstraps. The reported percentage denotes the proportion of vertices in an ROI classified consistently as either in-phase or anti-phase across all 500 resamplings of the data.*

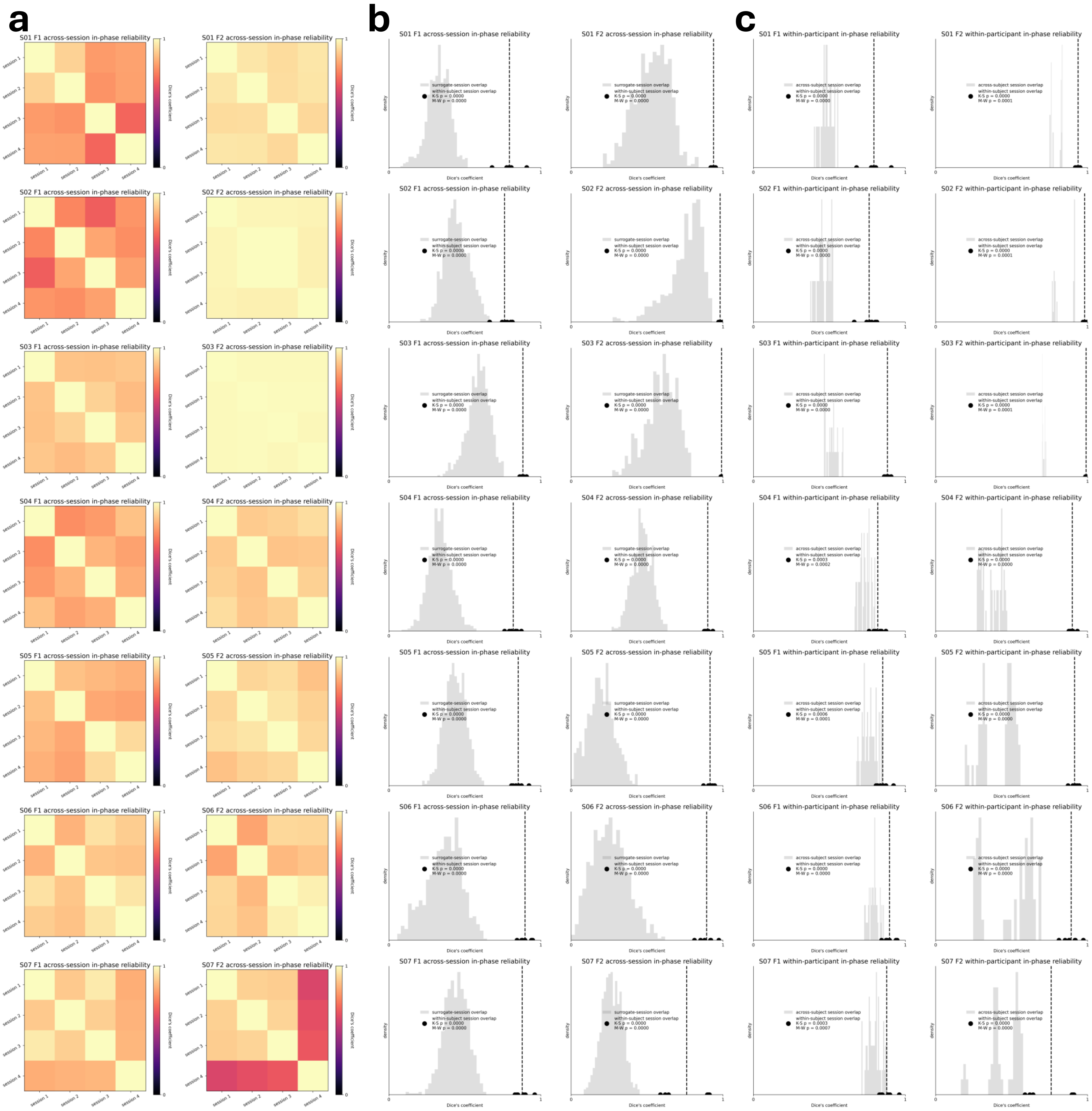
a
b
c
S01 F1 across-session in-phase reliability
S01 F2 across-session in-phase reliability
S01 F1 within-participant in-phase reliability
S01 F2 within-participant in-phase reliability
S02 F1 across-session in-phase reliability
S02 F2 across-session in-phase reliability
S02 F1 within-participant in-phase reliability
S02 F2 within-participant in-phase reliability
S03 F1 across-session in-phase reliability
S03 F2 across-session in-phase reliability
S03 F1 within-participant in-phase reliability
S03 F2 within-participant in-phase reliability
S04 F1 across-session in-phase reliability
S04 F2 across-session in-phase reliability
S04 F1 within-participant in-phase reliability
S04 F2 within-participant in-phase reliability
S05 F1 across-session in-phase reliability
S05 F2 across-session in-phase reliability
S05 F1 within-participant in-phase reliability
S05 F2 within-participant in-phase reliability
S06 F1 across-session in-phase reliability
S06 F2 across-session in-phase reliability
S06 F1 within-participant in-phase reliability
S06 F2 within-participant in-phase reliability
S07 F1 across-session in-phase reliability
S07 F2 across-session in-phase reliability
S07 F1 within-participant in-phase reliability
S07 F2 within-participant in-phase reliability
session 1
session 2
session 3
session 4
Dice's coefficient
density
surrogate-session overlap
within-subject session overlap
across-subject session overlap

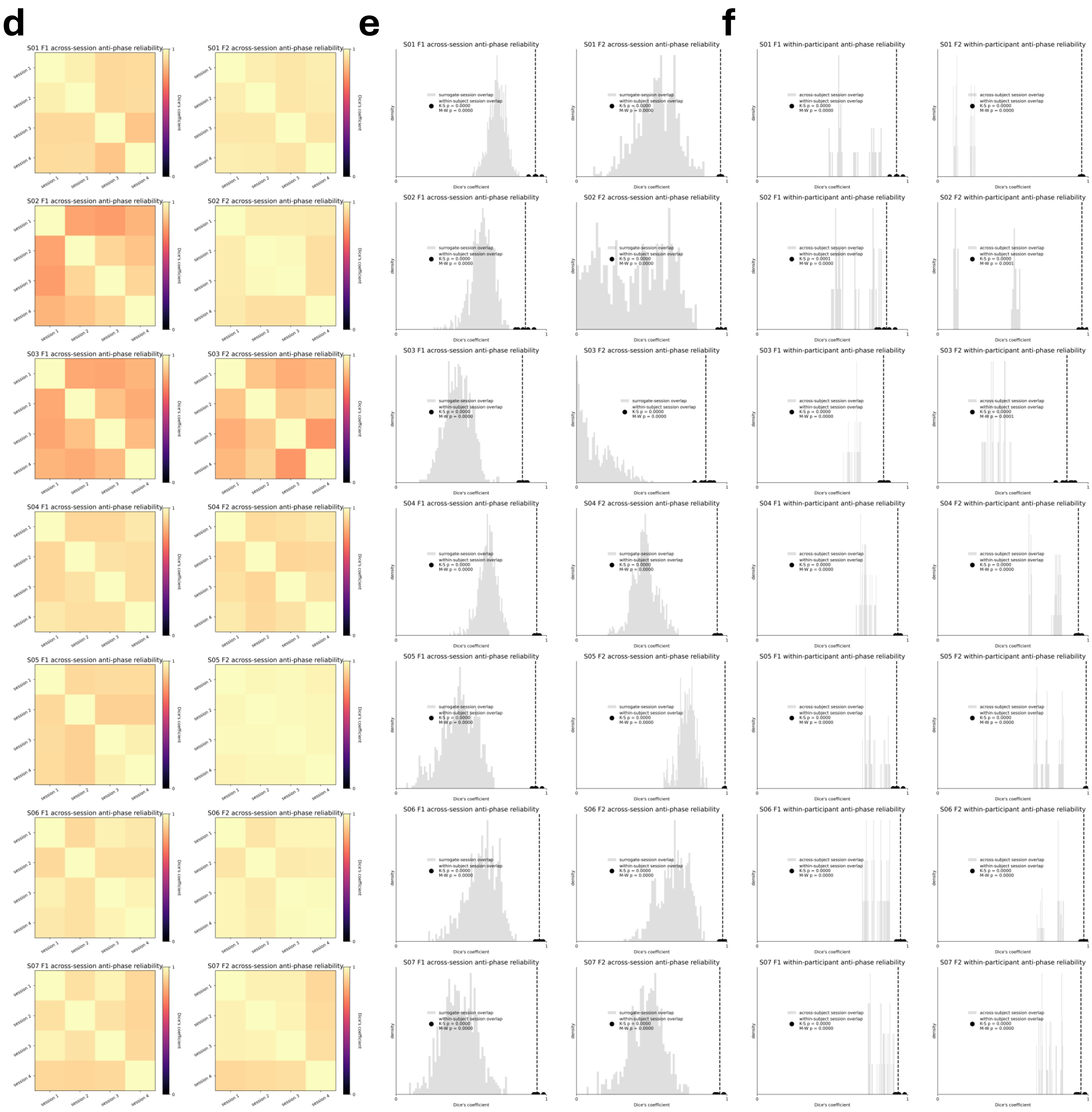


***Supplementary Figure 6: Reliability of signed phase maps across sessions and within participants***

***(a)*** *Overlap (Dice's coefficient) between in-phase synchronized vertices estimated from each experimental session (4 sessions; 4 runs of data each) for each participant (rows) at F1 (left) and F2 (right).* ***(b)*** *Surrogate map test to determine the significance of between session overlap of in-phase synchronized vertices. Data in gray shows the distribution of surrogate dice's coefficient values generated by computing the overlap between session signed phase maps with shuffled signed phase maps that retained the same spatial autocorrelation structure and anatomical distribution as the original F1 (left) and F2 (right) session signed phase maps. Black data points represent the between session attentional modulation map correlations within participants. Dashed vertical lines indicate the mean value of between session attentional map correlations. Reliability of in-phase synchronization was tested by comparing the between session Dice's coefficient values against the surrogate distribution for each participant (Kolmogorov-Smirnov and Mann-Whitney U $p_{one\text{-}tailed} < 0.05$ for 7/7 participants at F1 and F2). This test was repeated for the between session Dice's coefficient values and surrogate distributions aggregated across all participants (Kolmogorov-Smirnov $p_{F1\ one\text{-}tailed} = 1.07 \times 10^{-67}$; $p_{F2\ one\text{-}tailed} = 6.22 \times 10^{-39}$; Mann-Whitney U $p_{F1\ one\text{-}tailed} = 3.13 \times 10^{-29}$; $p_{F2\ one\text{-}tailed} = 3.23 \times 10^{-27}$).* ***(c)*** *Between participant session signed phase map test to examine the uniqueness of the functional topography of within-participant in-phase synchronized vertices. Data in gray shows the distribution of between-participant session signed phase map overlap values. Data representation follows the same convention as in (b). Individuality of in-phase signed maps was tested by comparing the within-participant between session overlap against the between-participant between-session overlap (Kolmogorov-Smirnov $p_{one\text{-}tailed} < 0.05$ for 7/7 participants at F1 and F2). This test was repeated for the within-participant between session Dice's coefficient values and between-participant between session Dice's coefficient values aggregated across all participants (Kolmogorov-Smirnov $p_{F1\ one\text{-}tailed} = 1.54 \times 10^{-13}$; $p_{F2\ one\text{-}tailed} = 2.51 \times 10^{-29}$; Mann-Whitney U $p_{F1\ one\text{-}tailed} = 2.97 \times 10^{-15}$; $p_{F2\ one\text{-}tailed} = 1.07 \times 10^{-21}$).* ***(d)*** *Overlap (Dice's coefficient) between anti-phase synchronized vertices estimated from each experimental session (4 sessions; 4 runs of data each) for each participant (rows) at F1 (left) and F2 (right).* ***(e)*** *Surrogate map test to determine the significance of between session overlap of anti-phase synchronized vertices. Representation of the data follows the same convention as in (b). Kolmogorov-Smirnov and Mann-Whitney U $p_{one\text{-}tailed} < 0.05$ for 7/7 participants at F1 and F2. Kolmogorov-Smirnov of aggregated overlap distributions ($p_{F1\ one\text{-}tailed} = 7.13 \times 10^{-97}$; $p_{F2\ one\text{-}tailed} = 1.45 \times 10^{-61}$). Mann-Whitney U of aggregated overlap distributions ($p_{F1\ one\text{-}tailed} = 2.54 \times 10^{-29}$; $p_{F2\ one\text{-}tailed} = 3.36 \times 10^{-29}$).* ***(f)*** *Between participant session signed phase map test to examine the uniqueness of the functional topography of within-participant anti-phase synchronized vertices. Representation of the data follows the same convention as in (c). Kolmogorov-Smirnov and Mann-Whitney U $p_{one\text{-}tailed} < 0.05$ for 7/7 participants at F1 and F2. Kolmogorov-Smirnov of aggregated overlap distributions ($p_{F1\ one\text{-}tailed} = 3.97 \times 10^{-21}$; $p_{F2\ one\text{-}tailed} = 5.21 \times 10^{-42}$). Mann-Whitney U of aggregated overlap distributions ($p_{F1\ one\text{-}tailed} = 5.56 \times 10^{-22}$; $p_{F2\ one\text{-}tailed} = 3.52 \times 10^{-25}$).*

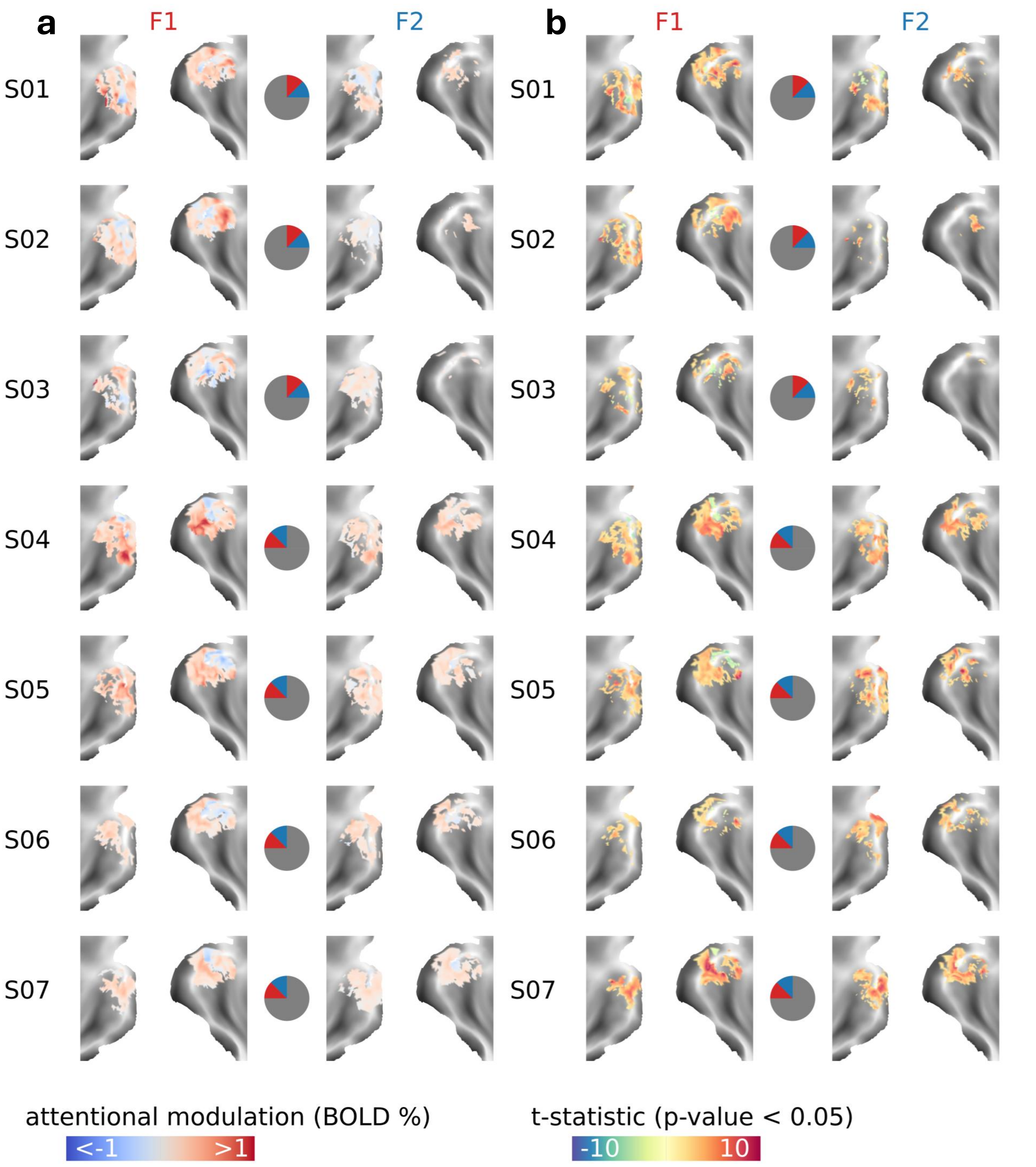


***Supplementary Figure 7: Individual participant attentional modulation maps***

***(a)*** *Participant attentional modulation maps at F1 (left) and F2 (right). The color bar denotes the extent of attentional modulation (F1: aF1-uF1; F2: aF2-uF2). The central diagram indicates the location of the oscillating wedge stimuli in the stimulated visual field quadrant (F1 in red; F2 in blue).* ***(b)*** *Participant t-statistic maps reflecting the significance of the effect of directed attention on oscillatory BOLD amplitude. Maps were generated by contrasting the amplitude of the BOLD oscillations at F1 during aF1 vs. uF1 (left) and at F2 during aF2 vs. uF2 (right). Contrast maps are thresholded at p < 0.05 (permutation test).*

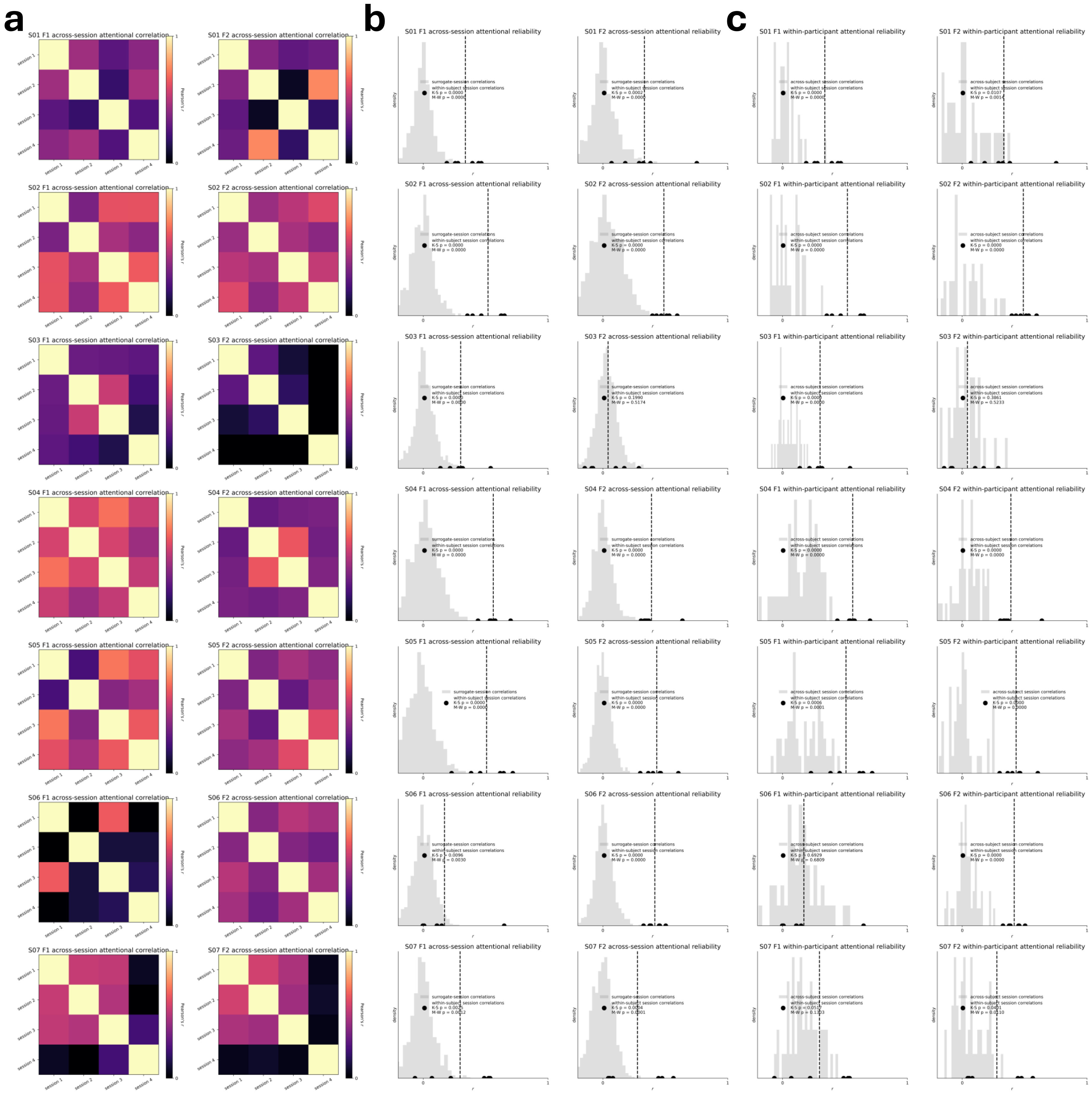


***Supplementary Figure 8: Reliability of attentional modulation maps across sessions and within participants***

*(**a**) Correlation (Pearson's r) between attentional modulation maps estimated from each experimental session (4 sessions; 4 runs of data each) for each participant (rows) at F1 (aF1-uF1; left) and F2 (aF2-uF2; right). (**b**) Surrogate map test to determine the significance of between session correlations. Data in gray shows the distribution of surrogate correlation values generated by correlating session attentional modulation maps with shuffled attentional modulation maps that retained the same spatial autocorrelation structure and anatomical distribution as the original F1 (left) and F2 (right) session attentional modulation maps. Black data points represent the between session attentional modulation map correlations within participants. Dashed vertical lines indicate the mean value of between session attentional map correlations. Reliability of attentional maps was tested by comparing the between session correlations against the surrogate distribution for each participant (Kolmogorov-Smirnov and Mann-Whitney U $p_{one\text{-}tailed} < 0.05$ for 7/7 participants at F1, and 6/7 participants at F2). This test was repeated for the between session correlations and surrogate distributions aggregated across all participants (Kolmogorov-Smirnov $p_{F1\ one\text{-}tailed} = 2.39x10^{-29}$; $p_{F2\ one\text{-}tailed} = 1.99x10^{-26}$; Mann-Whitney U $p_{F1\ one\text{-}tailed} = 1.41x10^{-23}$; $p_{F2\ one\text{-}tailed} = 5.37x10^{-20}$). (**c**) Between participant session attentional modulation map test to examine the uniqueness of the functional topography of within-participant attentional modulation maps. Data in gray shows the distribution of between-participant session attentional modulation map correlation values. Data representation follows the same convention as in (b). Individuality of attentional modulation maps was tested by comparing the within-participant between session correlations against the between-participant between-session correlations (Kolmogorov-Smirnov and Mann-Whitney U $p_{one\text{-}tailed} < 0.05$ for 5/7 participants at F1 and 6/7 participants at F2). This test was repeated for the within-participant between session correlations and between-participant between session correlations aggregated across all participants (Kolmogorov-Smirnov $p_{F1\ one\text{-}tailed} = 1.95x10^{-11}$; $p_{F2\ one\text{-}tailed} = 9.10x10^{-21}$ ; Mann-Whitney U $p_{F1\ one\text{-}tailed} = 5.76x10^{-13}$; $p_{F2\ one\text{-}tailed} = 3.82x10^{-16}$).*

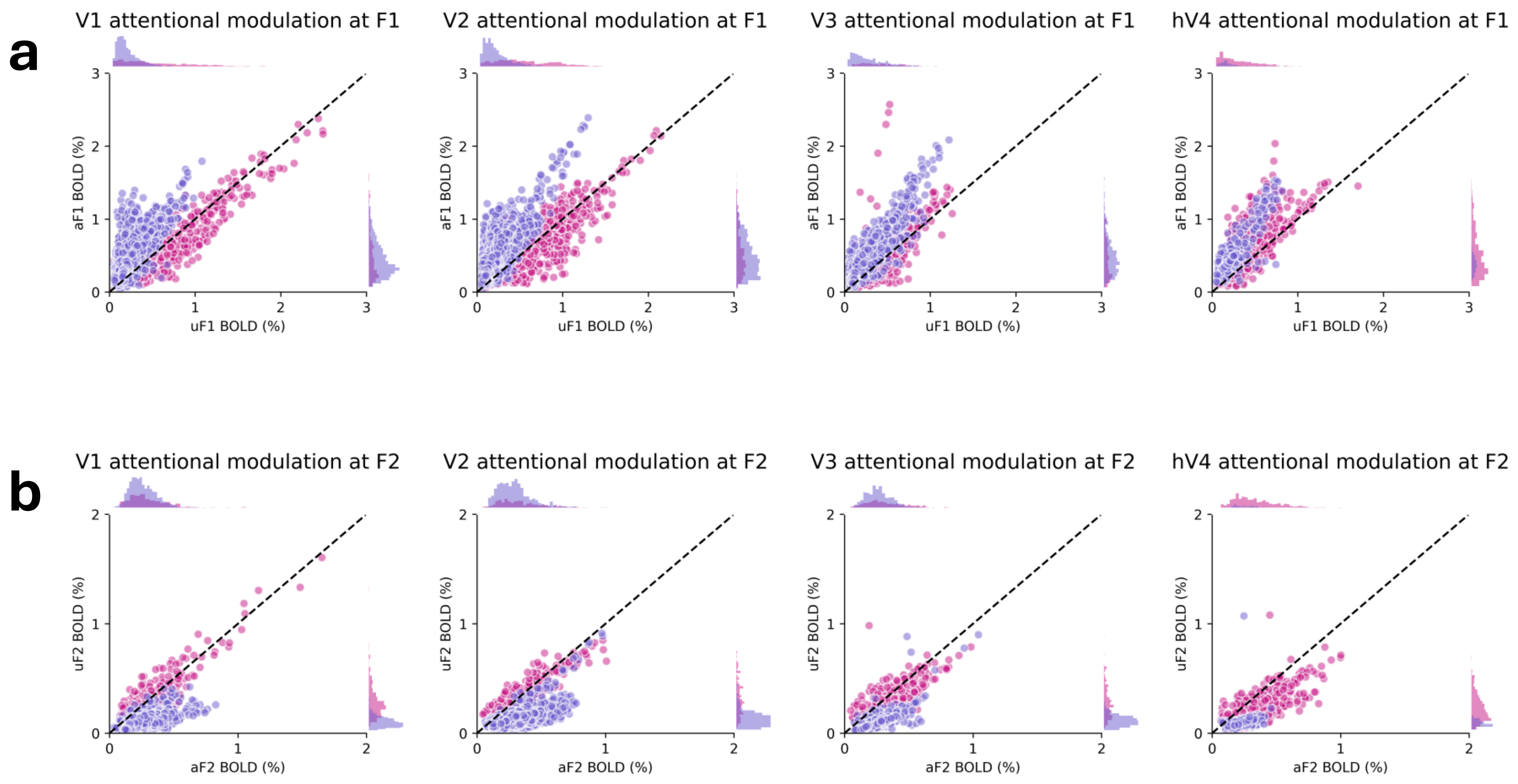


***Supplementary Figure 9: Attentional modulation of oscillatory BOLD amplitude of in-phase and anti-phase vertices in different visual ROIs***

***(a)*** *The relationship of aF1 (y-axis) to uF1 (x-axis) for visual ROI (columns) vertices of all participants for the in-phase (violet-red) and anti-phase (slate-blue) vertices. Dots along the diagonal indicate no difference in BOLD modulation between aF1 and uF1.* ***(b)*** *The relationship of aF2 (x-axis) to uF2 (y-axis). Representation of the data follows the same convention as in (a).*

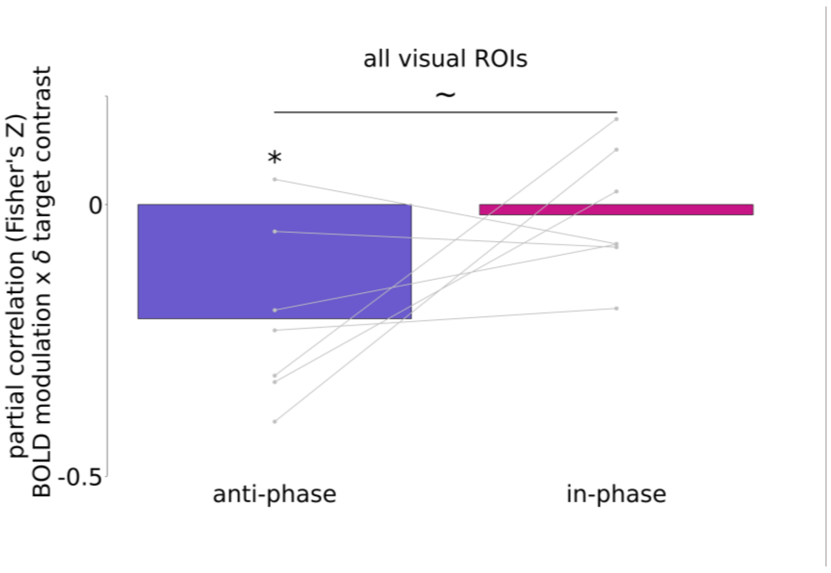


***Supplementary Figure 10: Attentional modulation of anti-phase BOLD oscillations predicts task performance above and beyond in-phase BOLD oscillations (partial correlation).***

*Run-by-run decreases in the fitted staircase slope (improvements in target sensitivity) relate to increases in the attentional modulation of anti-phase (slate-blue), but not in-phase (violet-red), oscillatory BOLD amplitudes when covarying for changes in the amplitude of the temporally opposing oscillatory population. Gray dots represent the participant Fisher's Z transformed spearman partial correlation value averaged across F1 and F2.*

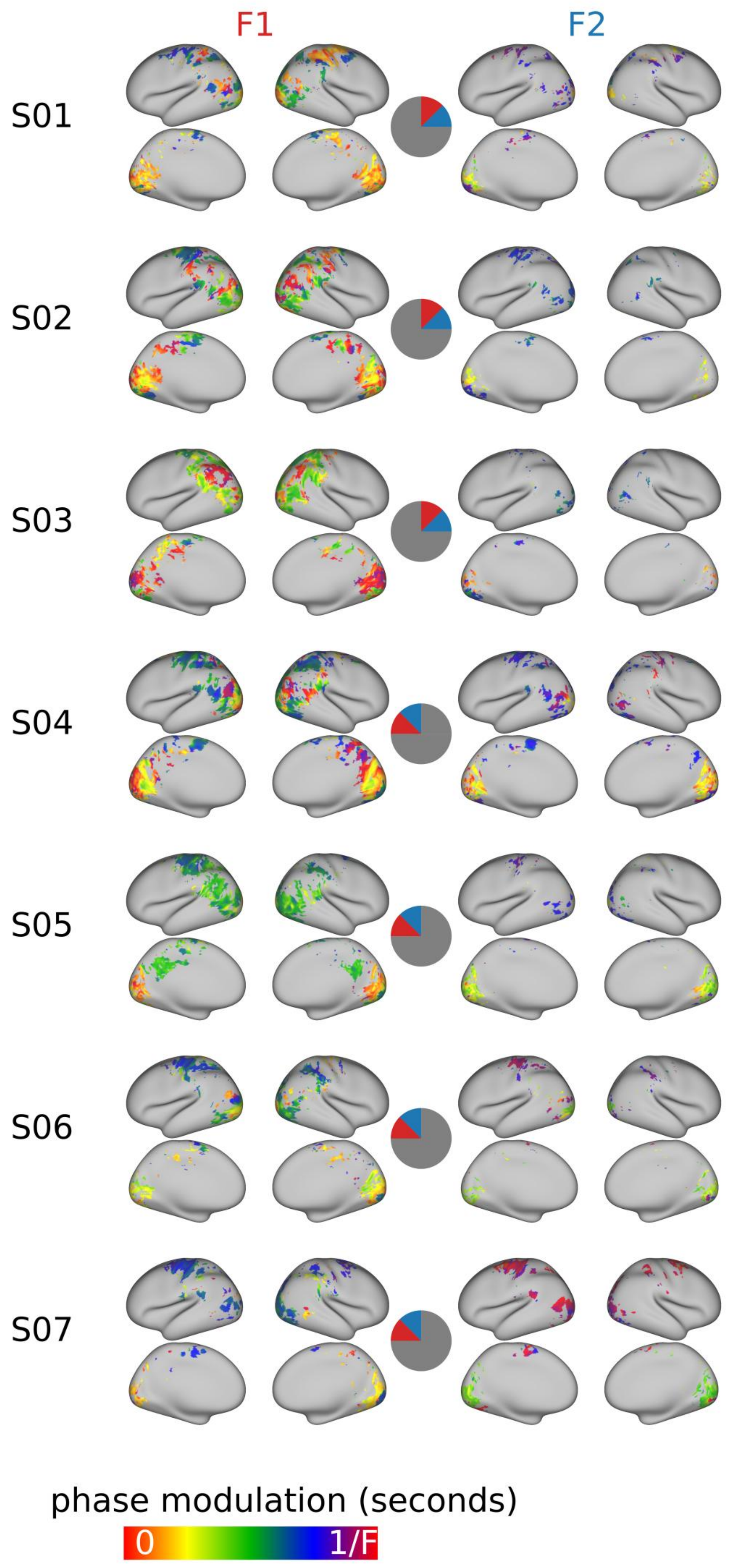


***Supplementary Figure 11: Thresholded participant phase maps (medial and lateral views)***

*Participant phase maps at F1 (left) and F2 (right). The color bar denotes the response phase scaled to the period of the examined frequency. The central diagram indicates the location of the oscillating wedge stimuli in the stimulated visual field quadrant for a given participant (F1 in red; F2 in blue).*

**Supplemental Table 1**. Test statistic = Wilcoxon signed rank tests with statistical null of no difference in the proportion of anti-phase vertices in an ROI across visual ROIs.

| Figure | ROI Comparison | Frequency | Test statistic | P-value |
|---|---|---|---|---|
| 3d | V1 vs. V2 | F1 | Rank = 3 | $p_{two\text{-}tailed} = 0.078$ |
| 3d | V1 vs. V3 | F1 | Rank = 7 | $p_{two\text{-}tailed} = 0.297$ |
| **3d** | **V1 vs. hV4** | **F1** | **Rank = 0** | $\mathbf{p_{two\text{-}tailed} = 0.0156}$ |
| 3d | V2 vs. V3 | F1 | Rank = 11 | $p_{two\text{-}tailed} = 0.688$ |
| **3d** | **V2 vs. hV4** | **F1** | **Rank = 0** | $\mathbf{p_{two\text{-}tailed} = 0.0156}$ |
| **3d** | **V3 vs. hV4** | **F1** | **Rank = 0** | $\mathbf{p_{two\text{-}tailed} = 0.0156}$ |
| 3d | V1 vs. V2 | F2 | Rank = 9 | $p_{two\text{-}tailed} = 0.467$ |
| 3d | V1 vs. V3 | F2 | Rank = 7 | $p_{two\text{-}tailed} = 0.297$ |
| **3d** | **V1 vs. hV4** | **F2** | **Rank = 1** | $\mathbf{p_{two\text{-}tailed} = 0.0313}$ |
| 3d | V2 vs. V3 | F2 | Rank = 9 | $p_{two\text{-}tailed} = 0.469$ |
| **3d** | **V2 vs. hV4** | **F2** | **Rank = 1** | $\mathbf{p_{two\text{-}tailed} = 0.0313}$ |
| **3d** | **V3 vs. hV4** | **F2** | **Rank = 2** | $\mathbf{p_{two\text{-}tailed} = 0.047}$ |

**Supplemental Table 2.** *S03 excluded due to lack of detectable anti-phase vertices in ROI. Test statistic = Wilcoxon signed rank tests with statistical null of no difference between the amplitudes of BOLD oscillations between the attended and unattended frequency.

| Figure | Phase population | Attended Frequency | Visual ROI | Test statistic | P-value |
|---|---|---|---|---|---|
| 4c | In-phase | F1 | V1 | Rank = 7 | $p_{two\text{-}tailed} = 0.297$ |
| 4c | In-phase | F1 | V2 | Rank = 7 | $p_{two\text{-}tailed} = 0.297$ |
| 4c | In-phase | F1 | V3 | Rank = 7 | $p_{two\text{-}tailed} = 0.297$ |
| **4c** | **In-phase** | **F1** | **hV4** | **Rank = 0** | $\mathbf{p_{two\text{-}tailed} = 0.0163}$ |
| **4c** | **Anti-phase** | **F1** | **V1** | **Rank = 0** | $\mathbf{p_{two\text{-}tailed} = 0.0163}$ |
| **4c** | **Anti-phase** | **F1** | **V2** | **Rank = 0** | $\mathbf{p_{two\text{-}tailed} = 0.0163}$ |
| **4c** | **Anti-phase** | **F1** | **V3** | **Rank = 0** | $\mathbf{p_{two\text{-}tailed} = 0.0163}$ |
| **4c** | **Anti-phase** | **F1** | **hV4** | **Rank = 0** | $\mathbf{p_{two\text{-}tailed} = 0.0163}$ |
| 4d | In-phase | F2 | V1 | Rank = 13 | $p_{two\text{-}tailed} = 0.938$ |
| 4d | In-phase | F2 | V2 | Rank = 9 | $p_{two\text{-}tailed} = 0.469$ |
| 4d | In-phase | F2 | V3 | Rank = 14 | $p_{two\text{-}tailed} = 1$ |
| **4d** | **In-phase** | **F2** | **hV4** | **Rank = 0** | $\mathbf{p_{two\text{-}tailed} = 0.0163}$ |
| **4d** | **Anti-phase** | **F2** | **V1** | **Rank = 0** | $\mathbf{p_{two\text{-}tailed} = 0.0163}$ |
| **4d** | **Anti-phase** | **F2** | **V2** | **Rank = 0** | $\mathbf{p_{two\text{-}tailed} = 0.0163}$ |
| **4d** | **Anti-phase*** | **F2** | **V3** | **Rank = 0** | $\mathbf{p_{two\text{-}tailed} = 0.0313}$ |
| **4d** | **Anti-phase*** | **F2** | **hV4** | **Rank = 0** | $\mathbf{p_{two\text{-}tailed} = 0.063}$ |
